# VISUALIZING THE NUCLEATING AND CAPPED STATES OF F-ACTIN BY Ca^2+^-GELSOLIN: SAXS DATA BASED STRUCTURES OF BINARY AND TERNARY COMPLEXES

**DOI:** 10.1101/2024.03.09.584204

**Authors:** Amin Sagar, Nagesh Peddada, Vikas Choudhary, Yawar Mir, Renu Garg, Ashish

## Abstract

Structural insight eludes on how full-length gelsolin depolymerizes and caps F-actin, while the same entity can nucleate polymerization of G-actins. Employing small angle X-ray scattering (SAXS) data analysis, we deciphered these two contrasting assemblies. Mixing Ca^2+^-gelsolin with F-actin in high salt F-buffer resulted in depolymerization of ordered F-actin rods to smaller sized species which became monodisperse upon dialysis with low salt G-buffer. These entities were the ternary (GA_2_) and binary (GA) complexes of gelsolin and actin with radius of gyration and maximum linear dimension of 4.55 and 4.68 nm, and 15 and 16 nm, respectively. In contrast, upon mixing G-actin with Ca^2+^-gelsolin in G-buffer, rapid association of higher order species started. Using size exclusion chromatography in-line with SAXS, we confirmed that initially GA and GA_2_ species are formed as seen upon depolymerization of F-actin, followed by dialysis. Interestingly, while GA_2_ could seed formation of native-like F-actin in both G- and F-buffer, GA failed in G-buffer. Thus, GA_2_ and GA are the central species formed via depolymerization or towards nucleation. SAXS profile referenced modeling revealed that: 1) in GA, actin is bound to the C-terminal half of gelsolin, and 2) in GA_2_, second actin binds to the open N-terminal half accompanied by dramatic rearrangements across gelsolin’s g1-g2 and g3-g4 linkers. Importantly, first structural insight is provided into the two probable models for GA_2_ with two actins in parallel, but differentially stacked: one in polymerization competent, and other in incompetent manner, suggesting latter to represent capped state along with the inert GA.

**TOC Abstract:** 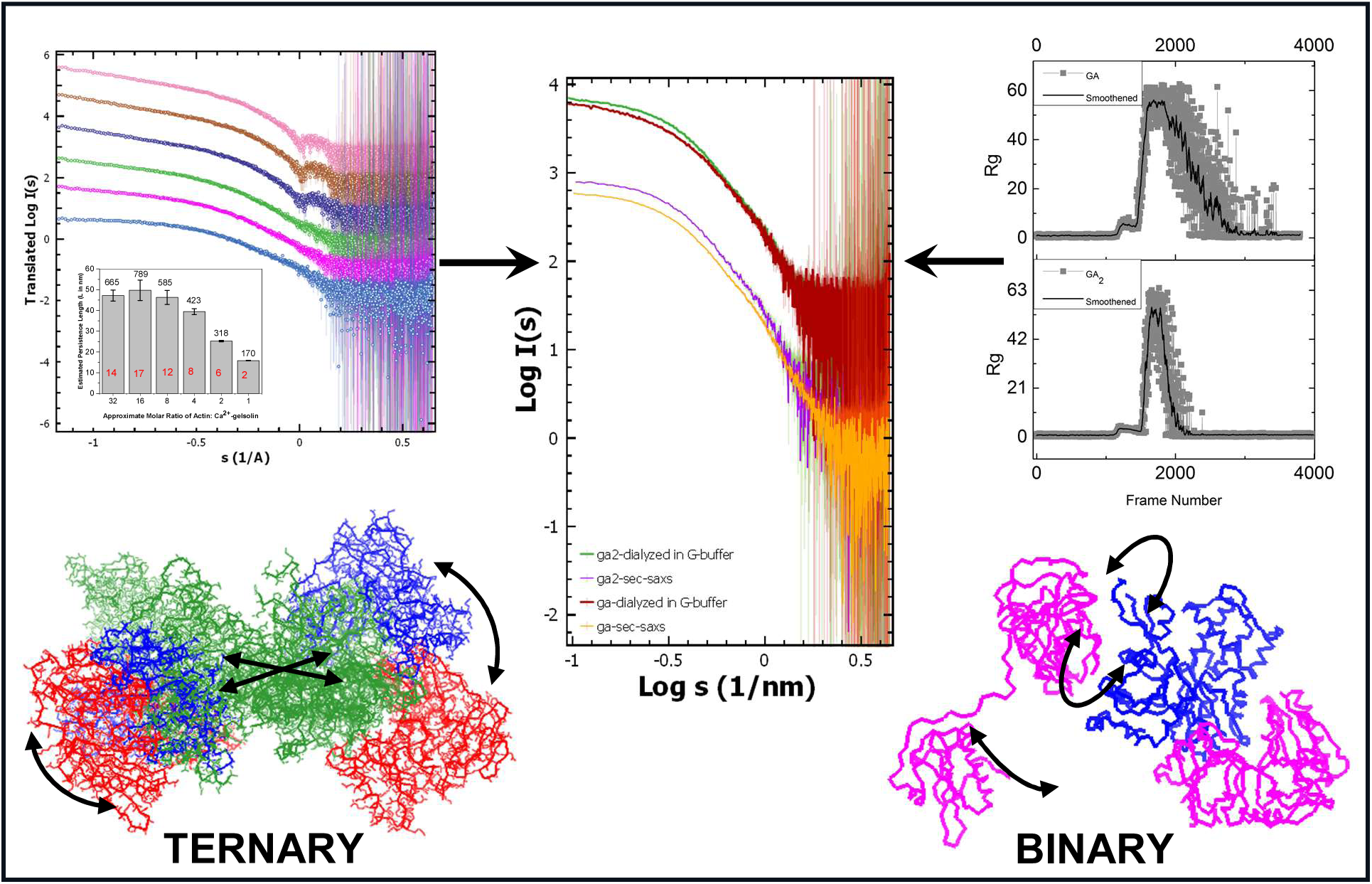

**Highlights:**

- Orderly decrement in the length of F-actin by Ca^2+^-gelsolin was tracked by SAXS.
- Residual re-association in 1:2 ratio in F-buffer was quenched by dialysis in G-buffer.
- Identical GA and GA_2_ entities formed upon mixing F- or G-actin with Ca^2+^-gelsolin.
- Models of nucleation competent, GA_2_ showed differential stacking of two G-actins.
- N-terminal half of gelsolin reposition as GA_2_ changes to or from GA, the capped state.

## INTRODUCTION

Gelsolin, a six domain (G1-G6), calcium sensitive protein is the only protein of the gelsolin/villin family which efficiently performs two seemingly opposing roles of actin nucleation as well as depolymerization [1]. A prerequisite for both processes is the Ca^2+^ ion binding induced metamorphosis of gelsolin from a compact inert shape to an extended and open active shape [2]. This conformational change is a three stage process as revealed by synchrotron radiolysis foot printing, tryptophan fluorescence and SAXS data analysis, which exposes the actin binding sites of gelsolin, effectively activating it for both severing and nucleation [2–4]. Apart from Ca^2+^, the severing activity of F-actin by gelsolin is also controlled by phosphatidylinositol 4, 5-bisphosphate (PIP2), Caspase-3, low pH and physiological temperature [5–8]. The binding with PIP2 sequesters gelsolin to the cell membrane and strongly inhibits the actin severing potency. On the other hand, Caspase-3 cleaves off the N-terminal half of gelsolin which interestingly possesses the ability to sever F-actin filaments in a calcium independent manner [9]. Recently, we showed that under Ca^2+^ free conditions, low pH and temperature close to 37°C can open up the G1 domain away from the remaining five enclosed domains being held together by Ca^2+^ sensitive C-terminal latch in gelsolin [7, 8]. The crystal structures of the different domains of gelsolin in complex with actin and mutagenesis studies have established the surfaces of contact between actin and gelsolin, and suggested mechanisms for severing and nucleation [10–12]. These structures show that the G1 and G4 domains of gelsolin bind between the subdomains 1 and 3 of actin in a similar conformation, while the G2 and G3 domains bind to the subdomains 2 and 1, respectively. The electron microscopy studies of gelsolin domains G2-G6 bound ADP-F-actin filaments suggest that gelsolin wraps around “the girth” of an actin filament such that the G2 domain binds between two actin monomers and C-terminal half (G4-G6) is located on the next actin protomer in the helical arrangement of actin molecules [13]. Upon activation by binding Ca^2+^ ions, full-length gelsolin is known to effectively depolymerize F-actin even in high ionic strength buffer and keep actin capped, thus eliminating actin from getting recommitted in F-actin formation. In contrast, some suggestions are there that Ca^2+^-activated gelsolin can bind up to two monomers of G-actin, and bring them together in space to initiate formation of F-actin [2, 14, 15].

Biochemical data concludes that a ternary complex of Ca^2+^-gelsolin and two actin units represent gelsolin’s capping and nucleation state, yet the plethora of structural studies available in the literature, a direct insight into the structure/global shape of this ternary complex eludes us. The complex of gelsolin with two actin molecules (GA_2_) is particularly important as it appears to be the common step in both severing/capping and nucleation. Previous attempts at obtaining diffraction competent crystals of GA_2_ have not been fruitful because of two main reasons: firstly, the full length gelsolin molecule in presence of Ca^2+^ ions is prone to proteolysis and gets digested into two fragments, during the time required for crystallization [6]. Secondly, in the “native-like” nucleating conditions, the system under study attains polydispersity due to the spontaneous self-association or gelsolin induced formation of actin filaments at concentrations typically required for getting diffraction quality crystals. Polydispersity and retention of native-like conditions remains a challenge for getting reliable electron density maps under cryo-conditions for electron microscopy without use of cross-linking agents or gold-tagged actins or small Fab arms. These limitations raised need for a nucleation incompetent mutant of one of the proteins which only provides skewed interpretation [16]. The structural assembly of this ternary complex has been further mystified by the reports showing contradictory evidence for the existence of an anti-parallel actin dimer during actin nucleation [17]. While the crosslinking studies suggest that the two actin molecules in a GA_2_ complex are highly mobile or in an anti-parallel conformation [17], ultracentrifugation studies of globular actin (G-actin) showed that actin dimers exist in negligible amounts compared to monomers and concluded that the anti-parallel actin dimers may be formed by the random collisions of monomers and these species “get trapped” in the presence of a crosslinking agent [18–20]. To further compound existing confusion, when the crystal structures of the two halves of gelsolin bound to actin (PDB IDs 1RGI and 1H1V) are opened together in any visualization program, they form a composite structure deceptively like the expected GA_2_ complex. In this study, we share how immaculately planned steady state and in-line size exclusion chromatography (SEC)-SAXS experiments allowed us to decipher the global shapes of these complexes at low concentrations in solution. The correlation of both approaches provided unambiguous conclusion that ternary (GA_2_) and binary (GA) complexes are central to both depolymerization or nucleation of F-actin by Ca^2+^-gelsolin in solution. The SAXS profiles provided structural constraints to model how extended Ca^2+^-gelsolin binds one actin, and then undergoes rearrangement to eventually hold two actin monomers or vice versa.

## Materials and Methods

### Gelsolin and Actin for Experiments

The cloning, expression and affinity purification of recombinant human gelsolin was performed as described previously [8, 21]. SAXS data was collected on purified gelsolin under EGTA and with free Ca^2+^ conditions which confirmed its foldedness and expected shape changes, respectively. The extraction and purification of actin was from chicken muscle was carried out as described by Spudich and coworkers with slight modifications as published before [8, 9, 22, 23]. SAXS data was collected on freshly purified G-actin sample to confirm its shape profile as published before [23].

### Preparation of GA_2_ and GA by mixing Ca^2+^-gelsolin with F-actin

For all the experiments gelsolin was activated by the addition of CaCl_2_ to a final concentration of 4 mM). The solutions of F-actin in F-Buffer (50 mM KCl, 50 mM Tris-HCl, pH 8.0, 5 mM MgCl_2_, 1 mM ATP, 0.1% 2-Mercaptoethanol) and calcium activated gelsolin in Buffer A (25 mM Tris-HCl, pH8, 45 mM NaCl, 1 mM EGTA) were mixed to a final concentration of 2-3 mg/ml with gelsolin to actin molar ratios of 1:1, 1:2, 1:4, 1:8, 1:16 and 1:32. [*Considering amount of EGTA in buffer, it is likely that about 3 mM of free Ca^2+^ ions were available to bind gelsolin and also influence chaotropic effects* [24]]. These mixtures were kept at room temperature (∼20°C) for 15 minutes before collecting SAXS data with gentle stirring. Importantly, before and after each sample, SAXS data was collected on mixtures of F-Buffer and Buffer A in the same ratio as the respective sample to be used for matched buffer subtraction. Additionally, one control experiment was done to prove the role of Ca^2+^ ions in depolymerization of F-actin by gelsolin. Like above, gelsolin to actin mixtures were made in similar molar ratios but with no addition of CaCl_2_ to gelsolin in Buffer A. As above, these mixtures were also kept at room temperature (∼20°C) for 15 minutes before collecting SAXS data with gentle stirring. As above, for each mixture, SAXS data was collected on mixtures of F-Buffer and Buffer A in the same ratio as the respective sample.

Additionally, F-actin and Ca^2+^-gelsolin were mixed in molar ratios of 1:1 and 1:2 and kept at room temperature for 30 minutes to allow complete depolymerization. Subsequently, to quench spontaneous filament formation and/or lower order association, the F-buffer was exchanged with low salt G-Buffer (2 mM Tris-Cl, pH 8.0, 0.2 mM ATP, 1 mM NaN_3_, 0.1 mM CaCl_2_ and 0.5 mM dithiothreitol, DTT) using Amicon protein concentrators with molecular mass cut-off of 10 kDa. Following buffer exchange with ∼20 times the sample volume, the samples were centrifuged at 13000 rpm for 15 minutes at room temperature with gentle stirring. The final concentration of the samples was ∼1.5 mg/ml. These samples were used for three sets of SAXS experiments: 1) As such, SAXS data collection with G-Buffer subtraction, 2) SAXS experiments were done with delay time of 15 minutes using aliquots taken out of mixing pool GA mixture in dialysis against G- and F-buffer, and 3) same experiments as before, but with pool mixture of GA_2_. These experiments were done to see if any association occurs in samples as a function of ionic strength of buffer and time. Bulk dialysis buffer in the set-ups were used as matched buffers.

### Preparation of GA_2_ and GA by mixing Ca^2+^-gelsolin with G-actin and using SEC-SAXS

Calcium activated gelsolin was mixed with G-actin in G-Buffer in ratios of 1:1 and 1:2 keeping the concentration of G-actin less than 0.6 mg/ml (in order to minimize spontaneous gelsolin-independent nucleation or concentration dependent self-association of G-actin [23]). The complex formation was incubated for 30 minutes at room temperature (∼20°C) and then concentrated using Amicon protein concentrators with cut-off of 10kDa. The samples with final protein concentration of ∼12 mg/ml (using absorbance at λ_280_ of gravimetrically diluted aliquots) were injected in a Superdex S200 column (pre-equilibrated with G-buffer) attached to the SAXS optics for data collection.

### Synchrotron SAXS data acquisition and processing

Synchrotron SAXS data was collected at P12 beamline at the Petra III storage ring at EMBL, Hamburg. All data was collected at 10°C using monochromatic X-rays and the sample to detector distance of 3.1 m. The scattered X-rays were recorded on Pilatus 2M detector, scaled, circular averaged about beam position, and respective buffer contribution was scaled and subtracted employing ATSAS pipeline. It should be noted that in all SAXS experiments injected samples were flowing at rate of about 30 µL/min to reduce any radiation induced anomalies in the exposed front of the sample. For each dataset, 20 frames of 45 milliseconds exposure time were collected. SAXS intensity profiles [I(s)] as a function of momentum transfer vector, s were obtained by circularly averaging intensity values recorded about the beam center. s is defined as relationship in Eq. 1:

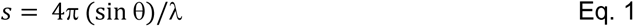

Units of s were in 1/nm. For each dataset, in automated manner, computed I(s) profiles of 20 frames were compared with each other and any profile with χ^2^ value below 0.95 or above 1.05 with previous frame were rejected from averaging. For each dataset, at least 12 similar SAXS profiles were averaged, and any set with less than 12 matching frames out of 20 collected frames was not considered. The averaged profile(s) of the matched buffer was subtracted from the averaged profile(s) of respective sample to obtain I(s) profile for the solute or protein molecules. All datasets were analyzed using ATSAS suite of programs v 3.0.3 [25]. Guinier analysis was done for buffer subtracted profiles considering globular or rod-like scattering shape of the particles, as required. Guinier analysis for globular and rod-like profiles provided radius of gyration (R_g_) and cross-section (R_c_), respectively along with extrapolated intensity value at s → 0 nm^-1^ (I_0_ values). Using relationship in Eq. 3, persistence length, L of the scattering species in

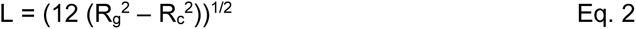

solution was estimated. Kratky plots and data comparisons were also generated/ done using inbuilt programs in the suite. The molecular masses of the particles were estimated from their SAXS profiles using Bayesian methods provided in the ATSAS suite. When required, indirect Fourier transformation of the SAXS intensity data yielded a pairwise distance distribution function of the inter-atomic vectors, P(r) as a function of vector length r, according to relationship in Eq. 3:

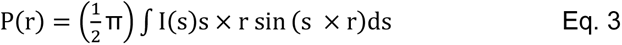

P(r) is the paired set of distances between all the electrons within the volume of the scattering particle. During this approximation, the probability of finding a vector equal to 0 nm and maximum linear dimension (D_max_) of the scattering particle was fixed to zero. The P(r) were calculated by the AUTOGNOM software available in the ATSAS suite of programs [25]. For the SEC-SAXS dataset processing, every 100^th^ frame was checked for identifying the frames with scattering intensity higher that the buffer scattering. The I_0_ and R_g_ for every 10^th^ frame of the selected set of frames was calculated in automated manner. These frames were checked for consistency in their auto estimated I_0_ and R_g_ values and then averaged in sets of 10 to improve the signal to noise ratio. The frames with minimal scattering closest to the elution peak were averaged to use for buffer subtraction. Key SAXS datasets used in this work are available in the SASBDB database and please see additional information [26].

### SAXS Data Steered Molecular Modeling

For GA and GA_2_ complexes, dummy residue chain-ensemble models were built using GASBOR program [27]. I(s) profiles were used as search reference, and P(r) analysis based deduced parameters were restraints for search dimensions. 1200 and 1600 dummy residues were used to model GA and GA_2_ complexes, respectively. For automated inertial axes alignment of dummy residue models and residue detail models were done using SUPCOMB program’s SASPy Plugin into PyMoL program [28]. Visual analysis, model building were done and images were made using PyMoL software [29]. The χ^2^ values between the theoretical SAXS profiles of different models and experimental SAXS data were computed and compared using CRYSOL program from ATSAS suite [30]. For some cases, residue detailed models were optimized to better resemble the information encoded in their SAXS data by searching low frequency normal mode analysis using SREFLEX program [31].

### SAXS based Structures of Ca^2+^-activated Full Length Gelsolin

Residue detail structure of Ca^2+^-activated gelsolin was generated by connecting the two crystal structures of N- and C-terminals of gelsolin (PDBs 1RGI and 1H1V) with modelled structure of the 70 odd residue g3-g4 linker [7, 9]. Earlier, we published that the six domains can be fitted inside volume and 3D shape profile of the dummy residue model of fully Ca^2+^-activated gelsolin, except the central “U-shaped” g3-g4 linker [2]. Different models of the linker were computed using PEPFOLD3 server (https://bioserv.rpbs.univ-paris-diderot.fr/services/PEP-FOLD3/) [32]. Shape profiles of the 100 computed models were first checked visually for the curved shape seen earlier via SAXS based modelling, and then calculated SAXS profiles of the selected curved models were compared with SAXS profile of the linker extracted from dummy atom model as done for another case [33]. The best fit was ligated theoretically with the crystal structures of N- and C-halves of gelsolin. The final model of full-length Ca^2+^-activated gelsolin was compared with its experimental SAXS data. The model was further improved by using experimental SAXS data and the SREFLEX program as detailed in the results section.

## RESULTS AND DISCUSSION

### G-actin, F-actin, and Ca^2+^-gelsolin for SAXS experiments

The SAXS profiles of freshly purified G-actin in G-buffer replenished with fresh ATP via dialysis, and after dialysis with F-buffer containing fresh ATP are presented in **Supplementary Figure S1**. Concentrations of G-actin and F-actin in the experiments were about 0.6 and 1.5 mg/ml, respectively. Peak profiles of their Kratky profiles confirmed expected foldedness or globular profiles of these protein molecules in respective buffers. Guinier analysis of the data profiles and their distance distribution profiles of the interatomic vectors for Ca^2+^-gelsolin, G-actin and F-actin are shown in **Supplementary Figure S1B, S1C and S1D**, respectively. As published before [2], the estimated P(r) profile for Ca^2+^-gelsolin showed an extended shape profile with D_max_ and R_g_ of 16 and 4.34 nm, respectively. Normalized Kratky profile for Ca^2+^-gelsolin had its peak maxima at values higher than sR_g_ close to 1.73 and had a plateau profile implying Gaussian-chain like disorderly behavior for the Ca^2+^-gelsolin molecules in solution. SAXS profile G-actin sample showed solution shape with D_max_ and R_g_ of 6.8 and 2.47 nm, respectively. Normalized Kratky profile with peak maxima close to sR_g_ ∼1.73 confirming a globular folded shape for G-actin molecules. These shape parameters were comparable to those published earlier by us [23]. Concentration of actin under G-buffer conditions were deliberately kept close to 0.6 mg/ml to quench any protein concentration induced self-association. SAXS profile of G-actin was compared with computed SAXS profiles of monomeric actin in different states of ATP binding, and results showed that our actin sample was resembling open shape seen in structure of ATP bound G-actin (PDB ID 1HLU) [34] which happens as a function of time with ATP getting hydrolyzed [23] (**Supplementary S1E**).

This G-actin was dialyzed against F-buffer, concentrated using membrane concentrators. Then protein concentration was estimated using band intensity for its monomeric state in Coomassie stained SDS-PAGE’s image vs. band intensities of albumin samples with known protein concentrations. The SAXS profile of F-actin thus prepared with concentration close to 1.5 mg/ml is shown in **Supplementary Figure S1A**, and its Guinier and distance distribution profiles are shown in **Supplementary Figure S1D**. The SAXS profile of F-actin showed the characteristic features of actin in filament order with a depression in intensity followed by peaks corresponding to real space distances of about 6.5 and 5.4 nm, respectively (**Figure 1A**, inset showing depression at s ∼0.1 Å^-1^ or 1 nm^-1^ by blue arrow). This supported that the actin units were assembled into the correct periodic helical arrangement in the filaments [35]. Similar features in SAXS profiles of F-actin have been reported earlier too [36–39]. We confirmed that dilution of F-actin in F-buffer retained the features albeit with the increase in noise at lower concentrations (**Supplementary Figure S2**). Using structure of F-actin as submitted in PDB ID 5OOE and extending the F-actin length using actin-actin relative positioning in space using Holmes Model of F-actin, we used CRYSOL program to compute and compare theoretical SAXS profiles of F-actin models of increasing length vs. experimental SAXS data (**Figure 1A**) [40, 41]. As can be seen in **Figure 1A**, increasing the length of the F-actin length resulted in computed SAXS profile which better matched with the experimental data. Computed χ^2^ values decreased from 7.7 to 1.5, as F-actin length (L) was increased from 170 to 500 Å with 5 to 18 actin units in helical order. Interestingly, the SAXS profile computed for the F-actin model with 18 actin units showed the depression and peak profiles as in experimental SAXS data suggesting similar sized particles and order exist in solution. Furthermore, the L value of this model of about 500 Å or 50 nm correlated well with the D_max_ value of 51 nm from Distance Distribution of SAXS data of F-actin presuming globular shape (**Supplementary Figure S1D**). Additionally, we did Guinier analysis and Distance Distribution estimation of F-actin sample using rod-like shape profile which provided cross-section radius and diameter of F-actin to be about 3.5 and 7 nm, respectively. The cross-sectional diameter of 7 nm matched well with dimensions seen from electron microscopy [42, 43]. Employing Eq. 2, we estimated the L value from SAXS data to be about 50 nm (as seen from modeling and distance distribution from globular profile). Importantly, R_c_ and L values estimated from dilution series of F-actin were about 3.5 to 3.7 and 48-50 nm, respectively. Retention of the distinct dip and peak profiles, and L values concluded that F-actin length and helical order is not affected by mere dilution of the protein sample.

**Figure 1.**
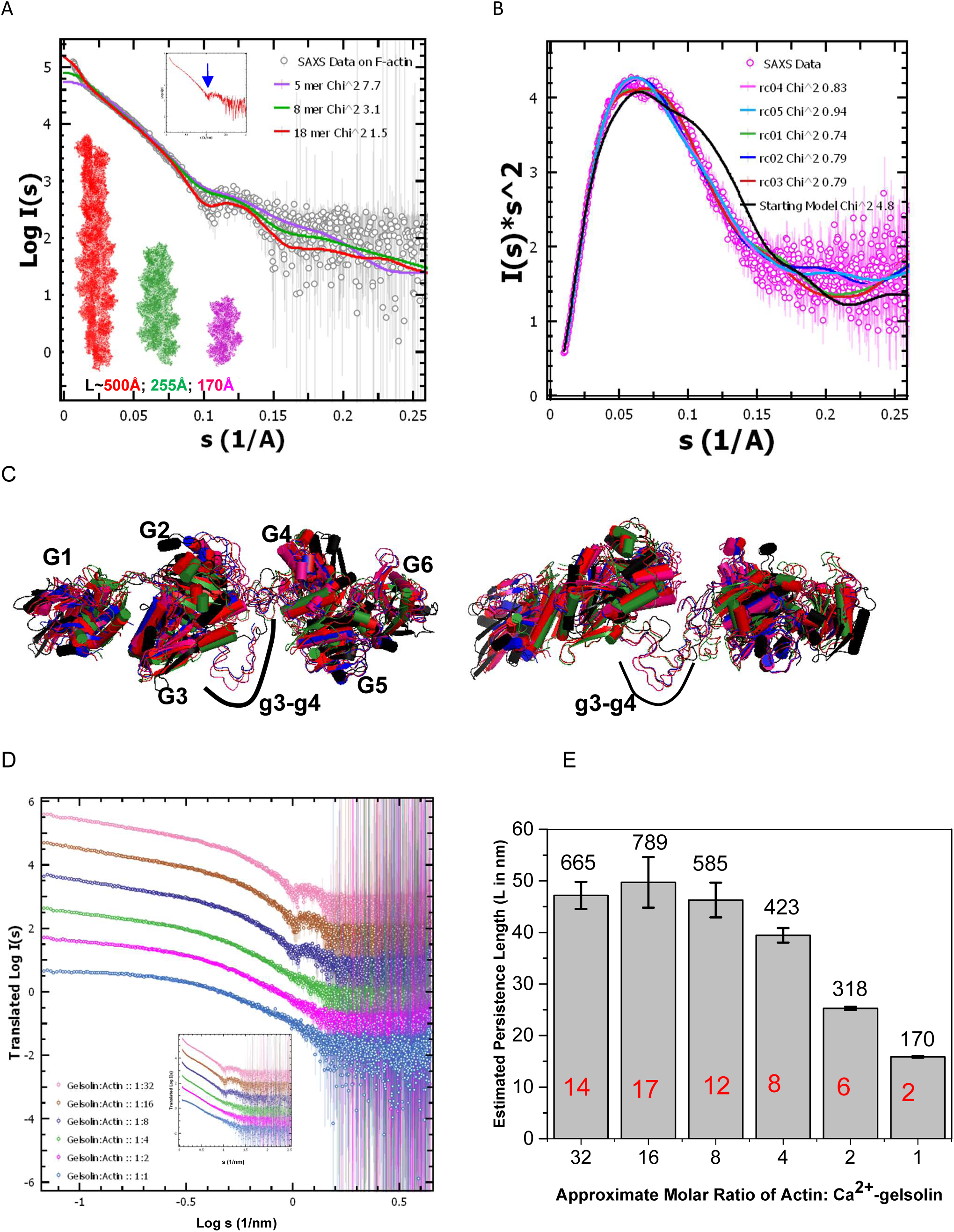
SAXS data-based characterization of F-actin and Ca^2+^-gelsolin used in this work. (**A**) SAXS data profile from F-actin sample (gray open circles) is plotted in Log-Linear mode. Inset shows the same plot without errors to pinpoint depression in the SAXS profile due to helical order in the filaments (blue arrow pointer). On the experimental dataset, theoretical SAXS profiles of models of F-actin filaments with persistence length of 500, 255 and 170 Å are shown in same colors as for models. In the legends, computed χ^2^ values vs. experimental data are mentioned. (**B**) The Kratky plots of experimental data from sample Ca^2+^-gelsolin (magenta open circles) are compared with same plots of SAXS data calculated from different models of full-length Ca^2+^-activated gelsolin (lines). Models are described in the main text and Supplementary Figure S3. Respective χ^2^ values to experimental data are also mentioned in the legends. (**C**) Six models of Ca^2+^-activated gelsolin (one initial in black color cartoon mode and five best results from normal mode analysis-based search) are overlaid here. Two rotated views show the spatial position of the six domains (labelled G1 through G6) and central g3-g4 linker. (D) SAXS profiles of different molar mixtures of actin in F-form and Ca2+-gelsolin in F-buffer are plotted here in Double Log mode. Ratios vary from 1:32 to 1:1 molar mixture of gelsolin to actin. Inset shows the same datasets in the Log-Linear mode. (**E**) This plot shows the decrement in the estimated value of persistence length of scattering species in different molar mixtures as gelsolin was increased. Numbers above histograms in black font mention the estimated molecular mass of the assemblies in SAXS data, and red fonts mention the estimation of actin molecules in the predominant scattering species.

### Open Structures of Ca^2+^-gelsolin within SAXS data-based constraints

Earlier, we published the SAXS data analysis and chain-ensemble dummy residue model of Ca^2+^-activated gelsolin [2]. When that model was compared with crystal structures of N- and C-halves of Ca^2+^-gelsolin, it showed that the two halves open-up with the central g3-g4 linker acting like a curved spacer. This model served as a basis to solve subsequent models of partially activated gelsolin [7, 8] and truncated forms of gelsolin [9, 44]. As of today, there is no residue detail model of Ca^2+^-gelsolin, and most prediction servers including DeepMind predict model of only inactive gelsolin with all six-domains compactly packed (as in PDB ID 1D0N chain A) [15]. SAXS profile of Ca^2+^-activated gelsolin are shown in **Supplementary Figures S1B and S3A**. As mentioned in the methods section, a model of central g3-g4 linker was generated and used to “stitch” together the two halves of Ca^2+^-gelsolin. Computed χ^2^ value of this model to the experimental SAXS data was 4.8 (**Supplementary Figure S3B**) (*In comparison, χ^2^ value computed using model of inactive gelsolin was ∼33*). Two rotated views of the residue detailed model representing Ca^2+^-gelsolin are shown in **Supplementary Figure S3C**. To solve structures of gelsolin which better fit the experimental SAXS data, we employed SREFLEX program which used the open model shown in **Supplementary Figure S3C**. This program employed normal mode analysis to search for better fitting models to the experimental SAXS profile. Computed SAXS profiles of the resultant models are compared to experimental data (**Supplementary Figure S3D**). The similarities between experimental data and profiles of searched models can be better appreciated in their comparative Kratky plots (**Figure 1B**). Interestingly, while the χ^2^ value of the initial model was 4.8, the SREFLEX program computed five models with χ^2^ values closer to unity and in the range of 0.74-0.94 (**Figure 1B**). Two rotated views of these five models are shown in **Figure 1C** which implies that the six domains can rotate about their spatial position mainly assisted by g1-g2 and g3-g4 linkers. This superimposition can represent structural ensemble which is accessible to Ca^2+^-activated gelsolin in solution. Apart from minor differences, the primary information one can conclude is that G1-G3 and G4-G6 halves of gelsolin are in the open state with their actin binding surfaces exposed. Below, we used the best-fitting model to generate models of gelsolin and actin complex.

### SAXS data profiles of F-actin depolymerized and capped by Ca^2+^-gelsolin

SAXS profiles of samples having Ca^2+^-gelsolin and F-actin mixed in molar ratios of 1:32, 1:16, 1:8, 1:4, 1:2 and 1:1 in F-buffer are plotted in Figure **1D**. It was expected that Ca^2+^-activated gelsolin will bind and depolymerize F-actin and cap the shorter actin filaments. Interestingly, the features characteristic of helical order of F-actin *i.e.* minima and maxima seen at 6.5 and 5.4 nm, gradually became less prominent pertaining to the loss of the periodic arrangement of the actin units and reduction in the filament length. At ratios of one gelsolin to four actin monomeric unit or more gelsolin, no minima or maxima characteristic of F-actin’s helical order could be seen indicating that all the actin filaments units had disintegrated into shorter entities. Parallel control experiments with inactive gelsolin (lacking free Ca^2+^ ions) showed that the rod-shape scattering, and depression-n-peak profiles were retained in mixtures having same ratios as in experiment shown in **Figure 1D**. The comparisons shown in **Supplementary Figure S4A** and **S4B** clearly support that Ca^2+^ ions are necessary for F-actin depolymerization activity of gelsolin. Furthermore, profile changes seen in **Figure 1D** are not due to mere addition of smaller sized Ca^2+^-gelsolin to rods of F-actin. Using the SAXS profiles presented from different mixtures in **Figure 1D**, cross-sectional Guinier approximations for rod-like shape was carried out to ascertain the relative ratio of the long *vs*. cross-sectional dimensions of the predominant scattering species in the F-actin/gelsolin mixtures. While linear profile of the data in the Guinier plots for rod-shape (ln I(s)*s vs. s^2^ plots) upheld a rod-like shape, “roll-over” trend of the plots at lower s values concluded that with increasing amount of Ca^2+^-gelsolin in mixtures, the long and cross-sectional dimensions of the scattering particles approached comparable dimensions (**Supplementary Figure S4B**).

Similarity in the slope of these plots and indirect Fourier transformation of the datasets for rod-shaped molecules confirmed that the cross-sectional diameter of scattering molecules remained comparable to that of F-actin alone *i.e.* about 7 nm at least till 1:8 molar ratio of Ca^2+^-gelsolin: F-actin mixture. Estimation of the L values for species in different Ca^2+^-gelsolin: F-actin mixtures showed that persistence length of particles decreased from ∼47 to 16 nm with increase in gelsolin in the mixtures (**Figure 1E**). Bayesian methods-based MOW analysis of the SAXS profiles suggested the average molecular masses of scattering species in solution (mentioned in black fonts above bars in **Figure 1E**). As expected, the molecular masses decreased from 660 kDa to about 170 kDa from 1:32 to 1:1 mixture. Presuming one gelsolin (∼80 kDa) is bound or capped to the depolymerized F-actin, number of actin units (∼42 kDa) were estimated (mentioned in red fonts in each bar of **Figure 1E**). For the datasets from 1:8 to 1:2 ratio mixtures, the estimated actin units in scattering species were higher than expected from complete depolymerization of F-actin by available Ca^2+^-gelsolin molecules. At this point, we realized that while it is expected that Ca^2+^-gelsolin can efficiently depolymerize and cap F-actin in higher ionic strength F-buffer, the same higher ionic strength buffer promotes F-actin formation. In other words, tread milling of actin molecules from capped state in lower order associations were getting recommitted to higher order associations in the F-buffer except for 1:1 mixture, and thus was causing artifacts in our SAXS profiles. To halt or quench this buffer ionic strength-based recommitment, for the 1:2 and 1:1 mixtures, we exchanged the buffer of the gelsolin-actin complexes formed in F-buffer into G-buffer as outlined in **Figure 2A** and the methods section.

**Figure 2.**
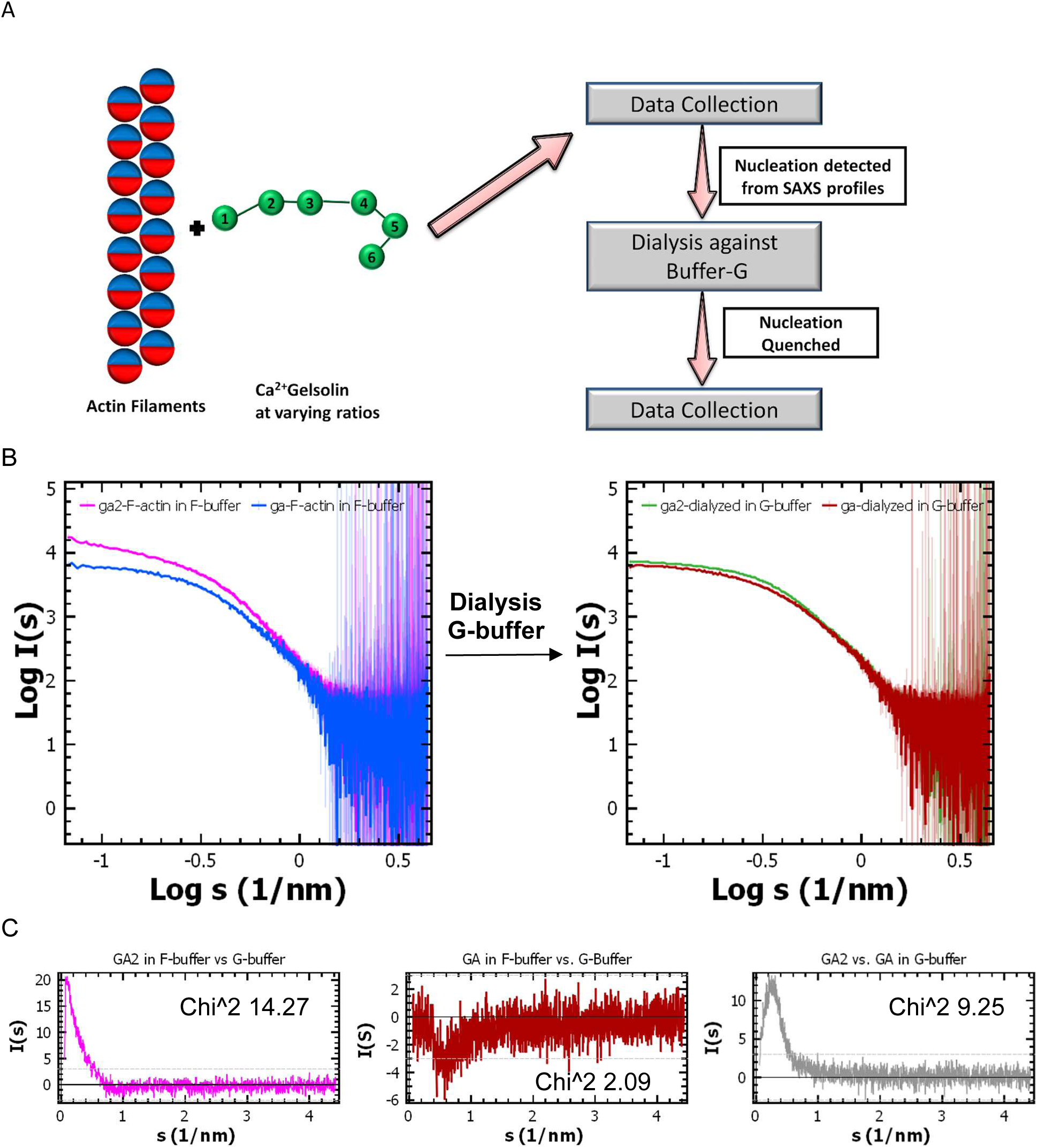
Results from dialyzing lower order ratios in G-buffer. (**A**) Schematic of the experimental set-up is shown here. Note that blue and red colors represent the bipolar ESP of actin units stacked in F-form. (**B**) Two panels show how the datasets for 1:2 and 1:1 mixture altered upon change from F- to G-buffer. (**C**) Comparison of the different datasets and the estimated χ^2^ value for the dataset pair are shown.

As shown in **Figure 2A**, the 1:2 and 1:1 mixture of Ca^2+^-gelsolin and F-actin prepared in F-buffer were dialyzed against G-buffer, and then SAXS data was again collected. The comparative SAXS profiles of the datasets in F- and G-buffer are presented in **Figure 2B**. Clearly, the upward trend of intensity values as s values decreased in the SAXS profile of 1:2 mixture in F-buffer disappeared when the mixture was in G-buffer. This clearly supported that lower ionic strength quenched low order association in the 1:2 mixture. Comparison of the datasets of 1:2 mixture in F- vs. G-buffer resulted in χ^2^ value of 14.27 (**Figure 2C left**). In contrast, for 1:1 mixture, there was little detectable change in the SAXS profiles from F- to G-buffer as seen from low χ^2^ value of 2.09 between the two datasets (**Figure 2C middle**). Thus, the depolymerization and capping was complete even in F-buffer for equimolar protein sample. Comparison of SAXS profiles of 1:1 and 1:2 mixtures in G-buffer resulted in χ^2^ value of 9.25 indicating that the shape information in their samples differed substantially (**Figure 2C right**).

### SAXS data profiles of G-actin units binding to Ca^2+^-gelsolin

To probe whether the lower order complexes of gelsolin: actin formed during nucleating conditions adopt similar shape as capped ones obtained from F-actin depolymerization, mixtures of G-actin and Ca^2+^-gelsolin were prepared in G-buffer, concentrated, and then were analyzed by SEC-SAXS (**Figure 3A**). Eluting buffer was also G-buffer and eluting solution from column was exposed to X-rays in an exposure time-period or a frame of 50 milliseconds in flow mode, and the R_g_ of the eluting molecules was plotted as a function of the frame (please see the methods section). One clear difference was seen in the elution profiles of the 1:1 and 1:2 molar mixtures of G-actin and Ca^2+^-gelsolin – while the 1:1 mixture eluted as a broad peak with an extended eluting “tail”, the 1:2 molar mixture showed a “tighter” elution profile (**Figure 3B**). This clearly indicated that the ternary complex is much more stable than the binary complex. The SAXS I(s) profiles of the frames inside the red-boxed zones being comparable in estimated R_g_ values were averaged and analyzed to ascertain the solution shape profiles of the predominant entity eluting from columns (shown in **Figure 3C**). The SAXS profiles from SEC-SAXS experiments were significantly lower in intensity values, and thus prior to comparison scale factors of 8.17 and 9.5 were applied for GA2 and GA, respectively (**Figure 3D**). Importantly, comparison of the SAXS profiles of 1:2 and 1:1 mixtures prepared via mixing Ca^2+^-gelsolin with F-actin (followed by dialysis in G-buffer) vs. G-actin (followed by SEC-SAXS in G-buffer) indicated χ^2^ values of 0.95 and 1.66, respectively concluding high similarities in the shape of entities formed via two protocols.

**Figure 3.**
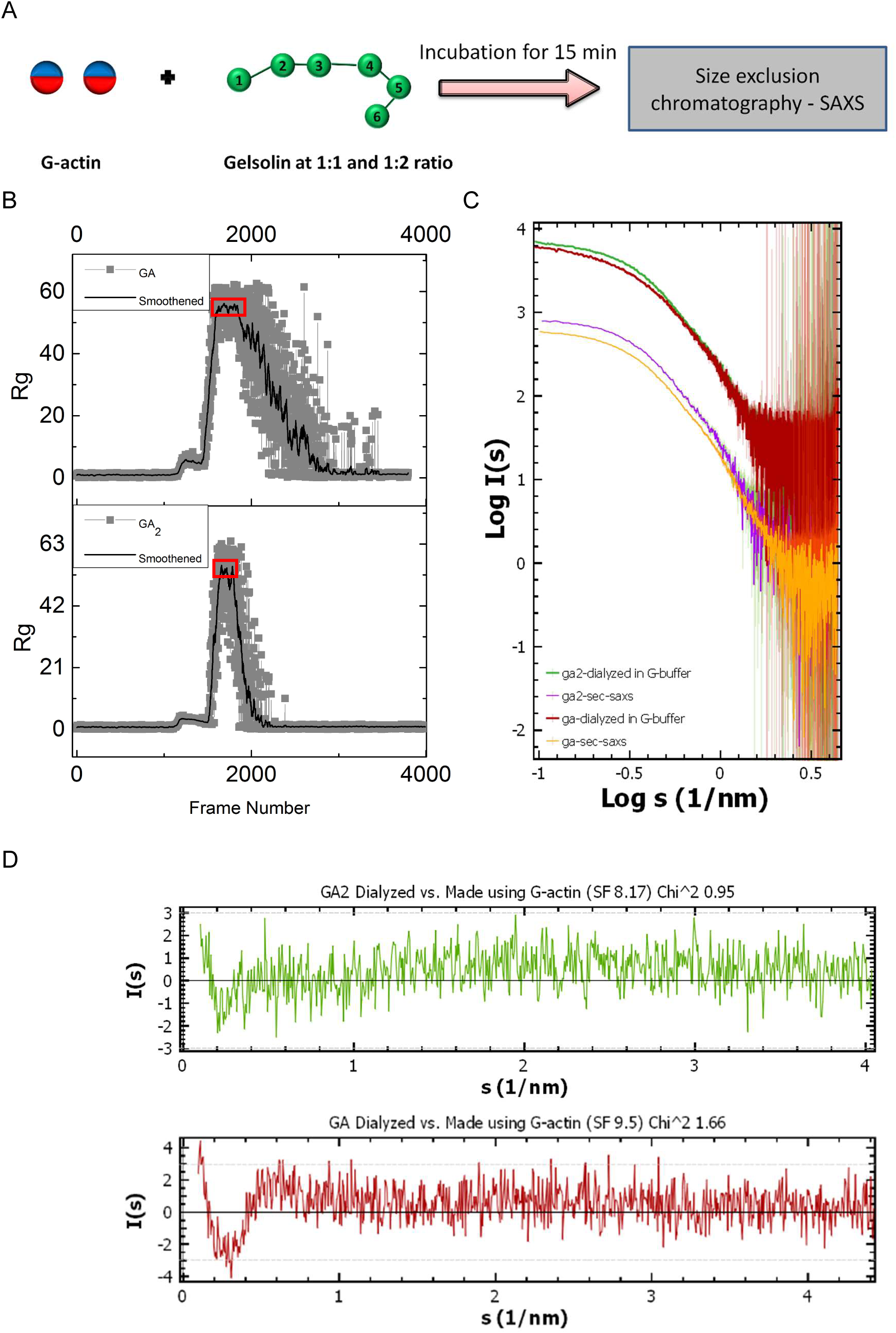
SEC-SAXS datasets obtained from mixtures of Ca^2+^-gelsolin and G-actin in G-buffer. (**A**) schematic showing the procedure used to prepare GA and GA2 complexes using G-actin and Ca^2+^ activated gelsolin and collect SEC-SAXS data. (**B**) Left: Plot of R_g_ as a function of the frame number in the SEC-SAXS experiments after incubating G-actin and gelsolin in 1:1 and 1:2 molar ratios are presented. The thick black line represents the smoothened curve obtained by adjacent averaging of 15 data points. The red box indicates the frames used for averaging and further processing. (**C**) The SAXS datasets of 1:2 and 1:1 molar mixtures prepared via F-actin depolymerization or complexation to G-actin are shown here in Double Log mode. (**D**) SAXS datasets of samples prepared in F-buffer and then dialyzed with G-buffer, and prepared in G-buffer followed by SEC are compared.

Preparative SEC experiments, prior to SEC-SAXS experiments shown above, also showed similar results that GA and GA2 prepared by mixing Ca^2+^-gelsolin with G-actin in G-buffer did not form large order associated species. Thus, a critical query was raised if these species in G-buffer could seed formation of F-actin like ordered assemblies. Earlier, like other researchers, we also followed F-actin formation by tracking increase in fluorescence by mixing pyrene labelled G-actin [8, 9]. Yet, those experiments did not confirm if the large order associations possessed the F-actin like orderliness seen by features in SAXS profiles? Or maybe the actin units required to form filament were all bound to Ca^2+^-gelsolin and thus limiting? Hence, the GA and GA2 mixtures were re-prepared as for SEC-SAXS experiments, and additional G-actin was added with final gelsolin: actin molar ratio to be about 1:16. Then, one part of mixtures was kept for dialysis in G-buffer and other in F-buffer having fresh ATP. At start and after about intervals of 15 minutes, a small fraction was taken out from dialysis cassette and SAXS data was collected. G- or F-buffer from dialysis was used for matched buffer profiles. For each set, about 2-3 minutes elapsed from sample extraction from dialysis cassette to data collection. Final SAXS profiles from the mixtures are presented in **Supplementary Figure S5A**. Results showed that in the SAXS experiments, scattering species associated having features like those for ordered F-actin in all the mixtures except for the GA + G-actin mixture in G-buffer. This was evident by the earlier seen depression and peak profile in the SAXS profiles in all three sets except for GA+G-actin in G-buffer. Automated estimation of the R_g_ values from the SAXS datasets showed that some increment in particle size occurs in the latter mixture, but the association rates were much faster and comparable for GA2+G-actin in G- or F-buffer, and for GA+G-actin in F-buffer (**Supplementary Figure S5B**). It was clear from these experiments that with the availability of additional G-actin units to stack as F-actin, GA2 is capable of nucleating ordered F-actin even in low ionic strength buffer. On the other hand, higher ionic strength F-buffer either promotes association of G-actin to gelsolin bound actin in GA sample, and/or leads to the formation of GA2 like shape which then nucleates formation of ordered F-actin.

Particle sizes and shape profiles of the predominant scattering species in the SAXS profiles of GA2 and GA mixture in G-buffer from SEC-SAXS are presented in **Supplementary Figure S6**. Guinier analysis presuming globular shape for 1:2 and 1:1 sample was 4.4 ±0.03 and 4.58 ±0.01 nm, respectively. Normalized Kratky plots showed peak profiles with maxima at sR_g_ value of 1.73 and slightly larger than sR_g_ value of 1.73 supporting globular and slightly lesser globular or disordered shape for the predominant particles in the two mixtures. Pair Distribution profiles of the interatomic vectors were estimated for both 1:2 and 1:1 mixture which provided D_max_ and R_g_ values for the predominant scattering shape in samples. They were 14.53 and 16 nm, and 4.53 and 4.67 nm for 1:2 and 1:1 mixture, respectively. Fit quality of the estimations can also be seen in the screenshots of PRIMUS Shape Wizard program in the **Supplementary Figure S6**. Additionally, by applying different methods, molecular weight analysis from SAXS datasets suggested that molecular masses were about 185 and 170 kDa with probability of estimation about 65% for 1:2 and 1:1 molar mixtures (**Supplementary Figure S6C**). Since, expected masses for the ternary and binary complex were about 160 and 120 kDa, higher numbers suggested that either there is some residual higher association in samples even in G-buffer and/or the partial specific volume of these species were higher than 0.74 cc/g which is considered for globular proteins [25, 45]. Please note that SAXS based shape parameters of Ca^2+^-gelsolin are presented in **Supplementary Figure S1B** which showed a disordered shape profile from normalized Kratky analysis of data and D_max_ and R_g_ of 16 and 4.34 nm, respectively. This suggested that addition of one and two molar equivalents of G-actin to Ca^2+^-gelsolin progressively increased globular nature of complexes, accompanied with increase in R_g_ and decrement in D_max_ value in 1:2 mixture.

### Structures of GA2 or Ternary Complex via SAXS-data based modeling

Using the structures of F-actin (PDB ID 5OOE), and N- and C-halves of Ca^2+^-activated gelsolin bound to one unit of actin each (PDB IDs 1RGI and 1H1V), we explored if shape information of ternary complex is present in the SAXS profile for 1:2 mixture obtained via depolymerization of F-actin by Ca^2+^-gelsolin followed by dialysis in G-buffer. Earlier, using SAXS data based modeling, we showed that Ca^2+^-activated gelsolin opens up so that sit across the “girth” of F-actin model and final binding would require movement of its G1 domain to bind its actin [2]. The protocol attempted in present work explored if the ternary complex generated via F-actin depolymerization still retained actin-actin contacts, while borrowing gelsolin-actin contacts from their co-crystal structures. As shown in **Figure 4**, actin units from the crystal structures 1RGI and 1H1V were superimposed over different combinations of adjacent actin units in the structure of F-actin. Options included actins in side-ways (models 1, 2, 6 and 7) as well as longitudinal contacts (models 3, 4 and 5). Additionally, placement of N- and C-terminal halves were swapped, as in models 1 vs. 2. Model 5 was ruled out as a possibility due to steric interactions which may arise between domains of gelsolin bound to actin as perceived from known crystal structures. Theoretical SAXS profiles of the six hypothetical models were computed and compared with the experimental SAXS data of 1:2 mixture of gelsolin and F-actin, then dialyzed in low ionic buffer. Different χ^2^ values and compared Kratky profiles for the six models are shown in **Figure 4**. Models 1 and 7, 2 and 6, and 3 and 4 are identical with computed χ^2^ values of 3.26, 1.82 and 3.09, respectively. Peak profile of all possible models had peak maxima close to that seen in the Kratky plot of the experimental data, but the upper peak widths were narrower and wider for models 1 or 7, and 3 or 4, respectively. Kratky profiles of models 2 or 6 had better fit to the experimental one and showed deviation mainly in the s range of 0.1-0.15 1/Å or 63 to 42 Å in real space. Latter dimensions mean that SAXS information on transfer vectors matched in large dimensions but disagree partially in the inner packing order or domain positioning as seen in the models 2 or 6 of the ternary complexes. The larger conclusion from this analysis is that models 2 or 6 with actin-actin side-ways contacts, best represent shape of the ternary complex seen in the experimental SAXS data via F-actin depolymerization (highlighted as red boxes in **Figure 4**).

**Figure 4.**
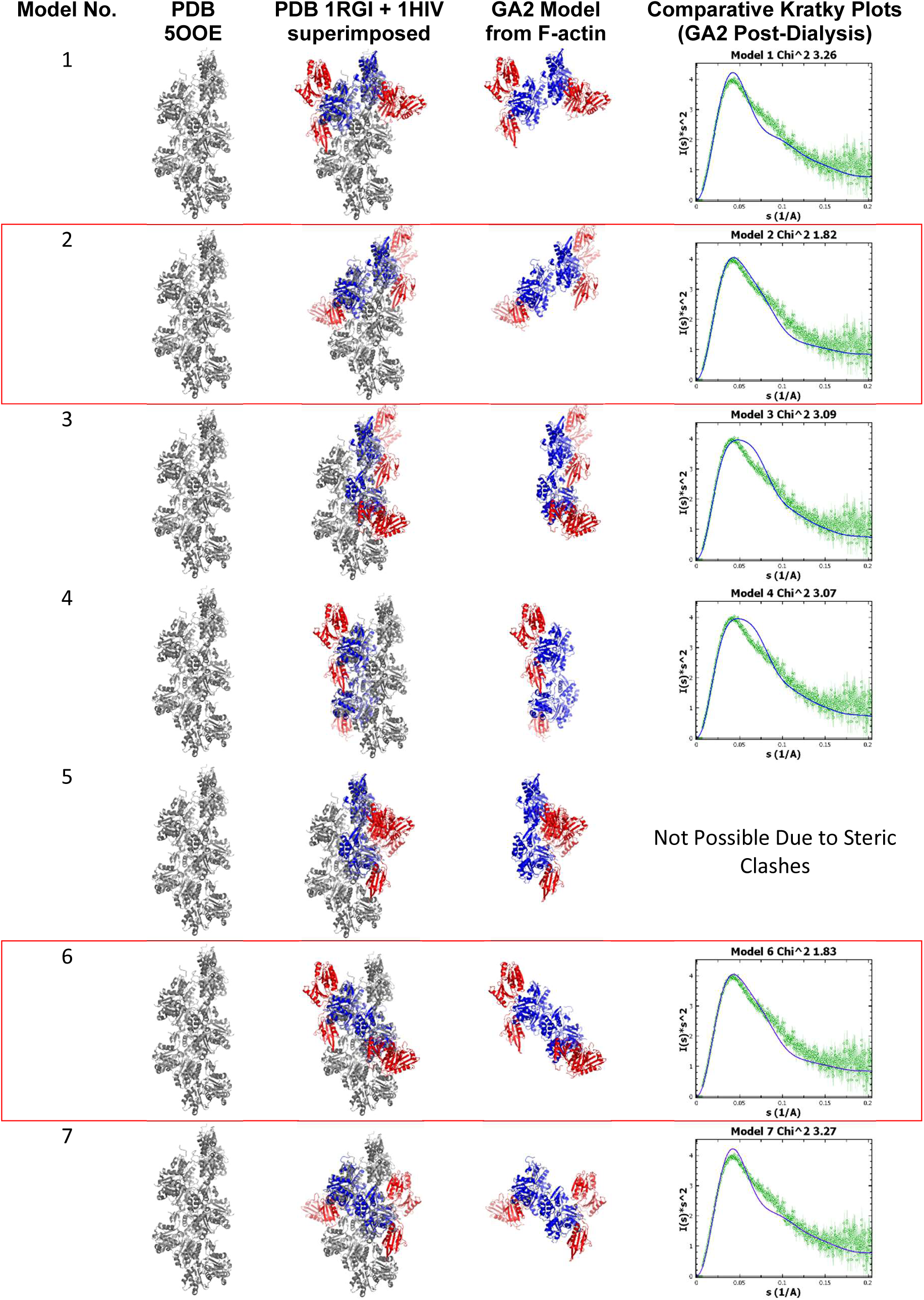
Different models of ternary complex considering adjacent chains of actin in the crystal structure of F-actin using actin and gelsolin contacts in crystal structures of N- and C-terminal halves bound to actin (PDB IDs 1RGI and 1H1V, respectively) are shown here. In columns third and fourth, red ribbons represent domains of gelsolin bound to their respective actins (blue ribbons) superimposed on actin chains in PDB 5OOE (gray ribbons). Right most column shows comparative Kratky plots of SAXS dataset from 1:2 mixture of gelsolin to F-form actin prepared in F-actin and dialyzed in G-buffer (green circles), are compared with calculated SAXS profile (blue line) from GA2 model shown in fourth column from left. Respective χ^2^ values of the comparison are also mentioned. Two rows i.e. for models 6 and 2 are boxed in red to highlight lowest χ^2^ values of all seven possibilities being considered here.

Using the SAXS data profile and deduced parameters for the 1:2 mixture of Ca^2+^-gelsolin with G-actin in G-buffer, we computed a dummy residue chain-ensemble model for the ternary complex (**Figure 5B** and **Supplementary Figure S7**). The χ^2^ value of the final SAXS profile of the dummy residue model was 0.76 to the experimental SAXS data (**Figure 5A**). Three rotated views of the solved model are shown in **Supplementary Figure S7A**. CRYSOL program computed different χ^2^ values for the seven ternary models generated using actin-actin contacts in F-actin structure vs. the SAXS data for 1:2 mixture from SEC-SAXS experiment are mentioned in **Figure 5A** along with their Kratky profiles. As earlier in the **Figure 4**, the χ^2^ values for models 1 or 7, 3 or 4 were higher than the values computed for models 2 or 6. Additionally, the peaks were wider and narrower for models 1 or 7 and 3 or 4 than the experimental data supporting disagreement in the shape information between theoretical models and experiment. As mentioned for earlier comparison, models 2 or 6 showed a χ^2^ value of 1.15 supporting that these models best represent the molecular shape of complex in the sample. One more model, named model 8, was generated by co-visualization of the two PDB structures of N- and C-halves of Ca^2+^-gelsolin bound to one actin molecule (PDBs 1RGI and 1H1V) (mentioned in the introduction). This model provided a χ^2^ value of 0.70 vs. the experimental SAXS profile. Considering χ^2^ value of unity to reflect best resemblance, structures in model 8 and 2 (or 6) best represent assembly of gelsolin and two actins in the ternary complex in solution (highlighted with black pointer arrows in **Figure 5A**).

**Figure 5.**
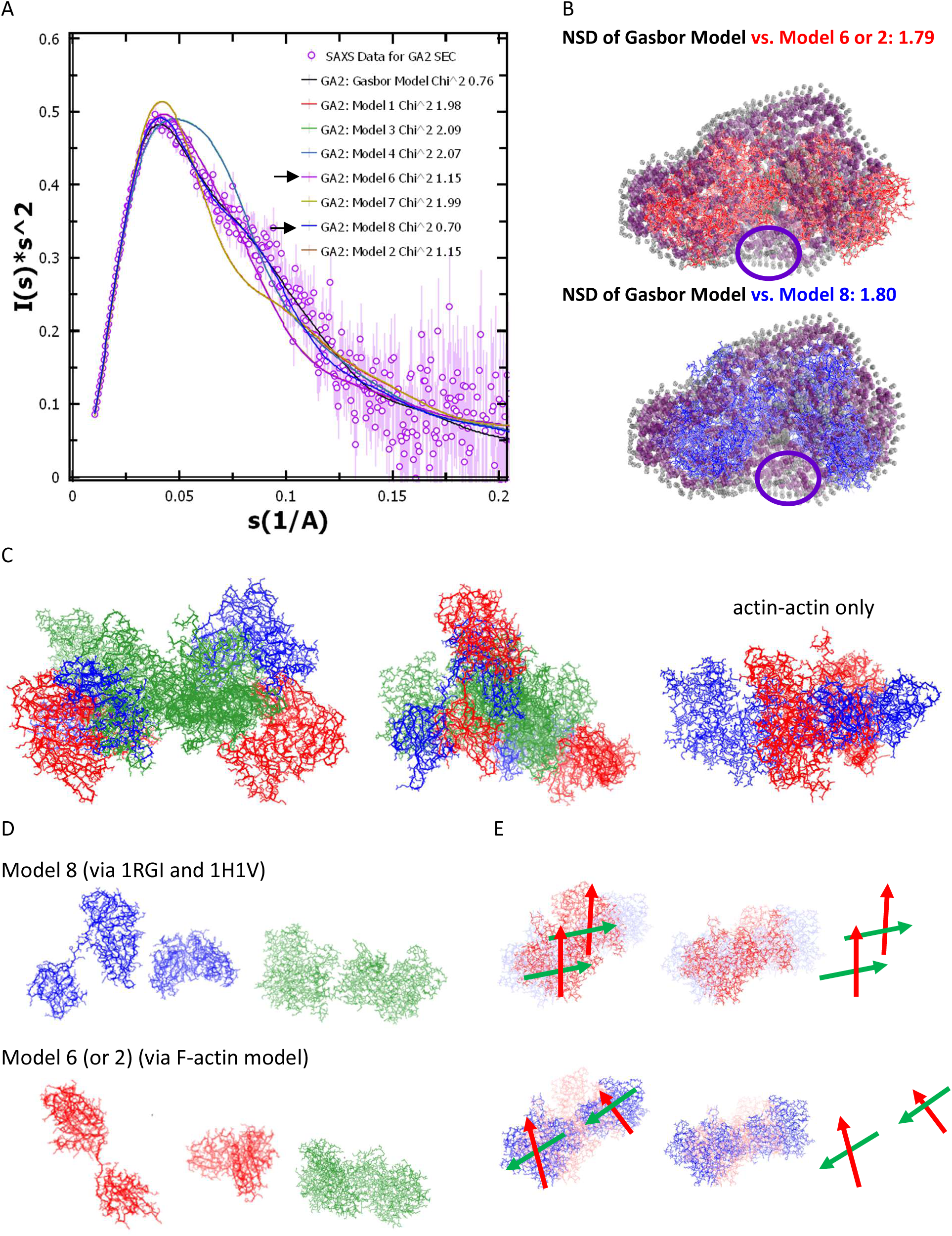
SAXS data-based models of ternary complex are presented here. (**A**) Comparative Kratky plots of different models of GA2 complex (lines) vs. SAXS dataset from SEC-SAXS of 1:2 mixture (violet circles) are plotted here. Respective line color and χ^2^ value for the considered model are mentioned in the legends. Two black arrows point towards the two sets which showed χ^2^ value closer to unity. (**B**) Inertial axes aligned superimpositions of the residue detail models of GA2 complex (red and blue lines) over the chain-ensemble dummy residue model (violet and gray cpk) are shown here. Gray cpk represents the hydration shell around the dummy residue (Asp; violet cpk). Normalized spatial disposition (NSD) values between the overlaid models are mentioned for each. The purple circle highlights unoccupied volume in the DR model where g3-g4 linker could have resided in the model 6 or 8. (**C**) Models 6 and 8 are aligned over their inertial axes (red and blue wireframes for the gelsolin domains bound to actins (green wireframes). The right most figure shows only the central actin units in the models 6 (red) and 8 (blue) of the GA2 complex. (**D**) Gelsolin domains as in model 8 and model 6 are shown in the left most panels followed by their respective actins (green wireframes). (E) Colors of central actins are now changed to red and blue for model 8 (upper) and 6 (lower), respectively. While actins in one model are in focus, the other have been faded for clarity. Two arrows red and green have been drawn to aid the reader on the direction of orientation of the actin chain, where red arrow runs from bottom to top (ATP binding site), and green arrow runs from domain residing DB loop to other end. The right most panel only shows the two sets of directional arrows.

By aligning inertial axes in automated manner, models 6 and 8 were independently aligned with dummy residue model (**Figure 5B** and **Supplementary** Figure 7B and C). The normalized spatial disposition (NSD) values were about 1.8 for both superimpositions with some penalty of unoccupied space in the dummy residue model (shown with purple circles in **Figure 5B**). It is relevant to note here that the g3-g4 linker connecting the two bound halves of gelsolin are not present in the residue detail models 6 or 8 but should have been modelled in the dummy residue model. This unoccupied volume in the dummy residue model highlighted may “house” the absent linker in models 8 (blue lines) or 6 (red lines). In the **Figure 5C**, left and mid panels show two views of the GA2 models superimposed on each other (RMSD value of all C^α^ atoms ∼ 27). Retaining the gelsolin-actin contacts as refined in the crystal structures, main differences were the spatial positioning of the G4-G6 halves and final placement of the G1 domain on opposite sides of the superimposition (with two actins shown in green lines). Only the two actins in the central part of the two models of ternary complexes are shown in the right panel of **Figure 5C** (blue for model 8 and red for model 6). Clearly, the two actins are differently oriented in the two models which agree with solution SAXS data. Additional views are presented in **Figure 5D** where the gelsolin halves and central actins are highlighted as in individual complex models. The superimpositions shown in **Figure 5C** right are reshown in **Figure 5E**, but with one set faded out for clarity. To guide eyes, we have added orientation vectors, where red and green arrows represent orientation and length from bottom of actin molecule towards its ATP binding pocket near open DB loop, and from DB loop half of actin to other half, respectively. This attempt for both models showed that though both the actins are parallel lengthwise, they are stacked face-to-face and end-to-end in models 8 and 6, respectively. More importantly, these models suggest that such orientational diversity may occur or simultaneously exist in gelsolin and actin ternary complexes in solution.

### Structures of GA or Binary Complex via SAXS data-based modeling

Four theoretical models were generated for the binary or GA complex of gelsolin and actin: <u>model 1</u>) actin bound to C-terminal half of best model of full-length Ca^2+^-activated gelsolin maintaining actin-G4-G6 contacts as seen in PDB ID 1H1V, <u>model 2</u>) actin bound to N-terminal half of full-length gelsolin maintaining actin-G2-G3 contacts as seen in PDB ID 1RGI, <u>model 3</u>) removing actin bound to N-terminal half of gelsolin from model 8 of GA2 complex, and <u>model 4</u>) removing actin bound to C-terminal half of gelsolin from model 8 of GA2 complex. CRYSOL program was employed to compute SAXS profiles of these models and compare them to the experimental SAXS profile (**Figure 6A**). The χ^2^ values of the models using structural model of full-length gelsolin are shown in box. Amongst the four options, the models with actin considered bound to C-terminal showed lower χ^2^ values compared to other ones *i.e.* model 1 and model 3 (highlighted by pointer arrows in **Figure 6A**).

**Figure 6.**
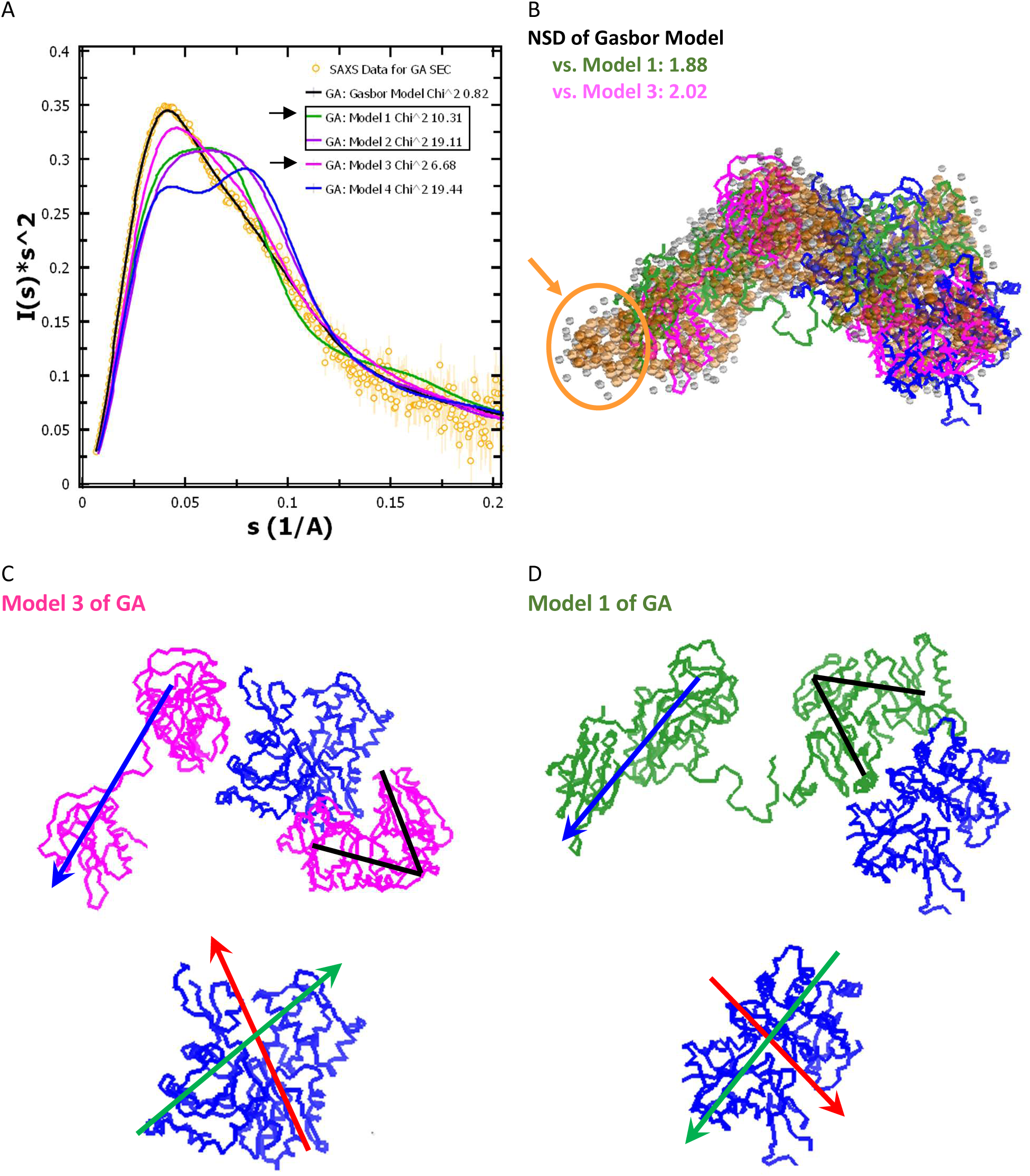
SAXS data-based models of binary complex are presented here. (**A**) Comparative Kratky plots of different models of GA complex (lines) vs. SAXS dataset from SEC-SAXS of 1:1 mixture (orange circles) are plotted here. Respective line color and χ^2^ value for the considered model are mentioned in the legends. Two black arrows point towards the two sets which showed χ^2^ value closer to unity. The values for the models generated using open structure gelsolin described in this work are in black box. (**B**) Inertial axes aligned superimpositions of the residue detail models of GA complex (gelsolin is shown in green and magenta for model 1 and 3, and actin chains are in blue) over the chain-ensemble dummy residue model (orange and gray cpk) are shown here. Gray cpk represents the hydration shell around the dummy residue (Asp; orange cpk). Normalized spatial disposition (NSD) values between the overlaid models are mentioned above. The orange circle highlights the disagreement between the spatial position of G1 domain in DR model vs. its position in the models 1 or 3. (**C**) Model 3 and (**D**) model 1 of GA have been presented as presented in the superimposition in the panel B. The directional arrows drawn to aid the reader are: blue for the G1-G3 domains, black V to show structure of G4-G6 domains, red and green arrows as in Figure 5D and 5E. *Please note the opposite orientation of V shape between model 3 vs. 1*. Figures in the lower row of this panel show the opposite orientation of actin chain in the two models.

Additionally, a dummy residue model of binary complex was generated using chain-ensemble model protocol, 1200 dummy residues and SAXS data-based parameters (**Supplementary Figure S8A**). The final χ^2^ value of this model vs. SAXS data was 0.82, much closer to unity than the variations computed for models 1 and 3 of GA complex. The Kratky plot of the dummy residue model showed a compact peak-n-shoulder profile like that for the experimental data (**Figure 6A**). Of the Kratky plots of the four models generated for the binary complex, only plot for model 3 showed a narrower peak with shoulder, while models 1 and 2 showed wider peak profiles than experimental data, and model 4 showed a two peak separated by a trough profile. Based on this comparison, model 3 possesses structural organization representing the molecules in the experiment, and model 1 could be the next probable one. The residue detail models (models 3 and 1) of the binary complex were superimposed over the dummy residue model by automatically aligning their inertial axes. In slight disagreement to the trend from the χ^2^ values, the computed NSD values of model 1 and model 3 vs. GASBOR program-based model were 1.88 and 2.02, respectively. In both superimpositions, there was an unoccupied volume in the dummy residue model close to where G1 domain was placed (please see pointer in **Figure 6B**), suggesting that G1 domain may be more open in the experimental data and G2-G6 domains of gelsolin bound to actin on the C-terminal side could be more compact than perceived in models 1 and 3. Actins in both models 1 and 3 are shown as blue ribbons, while domains of gelsolin are shown as green and magenta ribbons, respectively (**Figures 6B-D**). The big differences between models 3 and 1 are: 1) relatively opposite positioning of actin due to differential positioning of G4-G6 domain in structural model of the full-length gelsolin, and 2) compact G1-G3 half of gelsolin due to shorter conformation of g1-g2 linker in the model of the full-length gelsolin. As done earlier, we added orientation vectors in actin molecule which expectedly showed reverse profile between the two models suggesting that for both assemblies to co-exist or interconvert then central g3-g4 linker may do some serious reorientation with actin bound to C-terminal half which is a possibility considering the slightly disordered profile in Kratky plot of data.

## Conclusions

We provide new structural insights into the dynamic processes which are in synchronization during Ca^2+^-gelsolin mediated depolymerization, capping and nucleation of F-actin in solution. The open structure of Ca^2+^-activated gelsolin is known to bind one and two actin molecules, but their structure and their correlation with actin assembly regulation has been done indirectly through biochemical and fluorescence studies. In our attempts to form central players i.e. binary and ternary complexes of gelsolin and actin molecules, we considered both plausible routes: depolymerization of F-actin and sequential binding of G-actin molecules. Foremost, we characterized our reagents which confirmed globular shape of G-actin in G-buffer as published before by us [23], and the extended open shape of Ca^2+^-activated gelsolin [2]. As of now, no residue detail model of Ca^2+^-gelsolin is available, so we used our SAXS data profile to construct a model by stitching together known crystal structures of Ca^2+^-activated G1-G3 and G4-G6 domains, and a model of g3-g4 linker. Search for better representative models, this initial model led to closely resembling structural models of Ca^2+^-gelsolin which aids in visualizing ensemble of structures accessible to fully activated state of this protein. The SAXS profiles of F-actin in higher ionic strength F-buffer showed characteristic depression and peak profile which were independent of dilution confirming that F-actin molecules adopt rod shapes in solution with expected helical order. Overlaying theoretical SAXS profiles of F-actin assemblies on experimental data further confirmed the scattering shape, length and orderliness of F-actin molecules. Most researchers including us, earlier monitored F-actin assembly or depolymerization by tracking fluorescence values of pyrene-labelled actin which reflected association order, but did not provide further insights on spatial organization [7–9, 24]. Here, we tracked changes in the SAXS data profiles and deduced parameters as a function of increasing molar ratios of Ca^2+^-gelsolin to F-actin. Analysis provided clear evidence that with increase in gelsolin molecules, depolymerization and capping of F-actin is efficiently done. First, the orderliness of F-actin was lost, then the rod lengths were systematically reduced without much change in cross-sectional profile of rods. At lower ratios, the molecular mass estimations of depolymerized F-actin, capped by gelsolin suggested that either depolymerization was incomplete or some associations were still being induced by F-buffer. Upon dialyzing the 1:2 and 1:1 mixtures of Ca^2+^-gelsolin: (F-form of) actin vs. low ionic strength G-buffer, there was a clear decrease in particle size profile in 1:2 mixture, while the SAXS profiles of 1:1 mixture remained same. This confirmed that in F-buffer, some mid order associations were reforming in the 1:2 mixture, but complete depolymerization of F-actin was achieved by Ca^2+^-gelsolin at equimolar mixture.

Our SEC-SAXS data profiles unambiguously concluded that that the binary and ternary complexes of gelsolin and actin are formed with identical shape profiles, regardless of whether they are prepared by depolymerization and capping of F-actin, or by binding of G-actin units to gelsolin. SAXS profiles clearly confirmed that these entities are common to gelsolin mediated assembly or disassembly of actin. Additional experimental results showed that with addition of extra moles of G-actin and as a function of equilibration time, only ternary GA2 complex could enable F-actin formation even in G-buffer. This means that GA2 complexes serve as a ready template to bind additionally available actin units leading to F-actin assembly and may represent the seeding or nucleation state. Interestingly, dialysis against F-buffer allowed both binary and ternary state to nucleate F-actin suggesting that F-buffer either induces additional shape changes in GA making it closer to seeding GA2 and/or higher ionic strength promotes binding of actin units to actin bound to C-terminal of gelsolin in GA which may then associate to open G1-G3 half. Comparison of calculated and experimental SAXS profiles of GA2 suggested two types of assemblies are probable and may co-exist in solution. While, both GA2 models showed the two actins tightly held together by gelsolin domains, one model had edge-to-edge contacts and other had face-to-face associations. While the former agreed with associations seen in F-actin and thus may be considered as a ready seed for recruiting third actin to start nucleation, the latter had actins in nucleation incompetent assembly. Interestingly, this conclusion correlates with previously reported fluorescence based actin polymerization experiment for both ATP and ADP bound actin [46–48]. It was earlier concluded that GA2 may not be the true seed for polymerization and there was role of ATP or ADP bound state of actin as it affected closed vs. open shape of actin as free vs. bound to gelsolin [23]. Taken together, these results sum up that more factors could be involved in the inefficiency of GA_2_ to act as a seed for polymerization. Being aware of this and to minimize possibilities, we used fresh ATP replenishments in all our experiments, yet the half-life of bound ATP cannot be completely ruled out. Our models of GA2 suggest that this inefficiency may arise due to the non-polymerization competent arrangement of actin molecules in some assemblies of GA2.

Our nucleation competent model of GA2 is essentially structural insight into the mechanism of actin nucleation proposed by Burtnick *et.al.* [15] that the extension of the G3-G4 linker helps the domains G1 and G4 bind to two actin monomers and bring them closer into the correct conformation to act as a nucleus for filament formation. The nucleation incompetent model agrees with the information seen from the crystal structures of actin bound halves but implies that these gelsolin bound actin molecules eventually adopt a filament propitious conformation, but this probably happens at a later stage like the formation of GA3 or a larger entity. Alternatively, comparison of the two association patterns across actins in our models of the GA2 complex suggest that either both arrangements can occur simultaneously, or the two actins bind in one form, possibly face-to-face and then readjust to edge-to-edge format which can further bind actin units to GA3, GA4 leading to F-actin assembly. Another point worth discussing here is that despite the differences in the two actin-actin contact patterns, they were in parallel orientation in both of our models for GA2 complex. As mentioned before, crosslinking studies suggested that the two actin molecules in a GA2 complex are highly mobile and are in an anti-parallel conformation [17]. SDS-PAGE of different stages of gelsolin mediated nucleation showed two bands for actin dimers, upper dimer (UD) and lower dimer (LD). The LD was seen in the initial phases of actin polymerization and in paracrystals of chemically cross-linked actin [20]. LD was shown to migrate close to 86 kDa and was speculated to be having actins in anti-parallel orientation [49]. Using same cross-linking agents, F-actin resulted in UD which migrated close to 115 kDa and expected to have the actins in parallel disposition. Purified LD could bind Ca^2+^-gelsolin in 1:1 molar ratio, but it inhibited formation of F-actin [17]. Taken together, these results imply that the two actins in ternary complex with gelsolin are not fixed in the F-actin like orientation rather they adopt flexible orientations probably accompanied by motions in the gelsolin and ignoring ATP hydrolysis induced opening of bound actin(s). Whether the flexibility inside ternary complex can allow switching of actins from parallel to anti-parallel or vice versa appears bit farfetched? Particularly when the species formed in our SAXS data profiles either by depolymerization of F-actin or by adding G-actin are almost identical and the Kratky plots do not support higher disorder like unliganded Ca^2+^-gelsolin or for GA state. Realizing that our models of GA_2_ has actins in the parallel orientation, alternative explanation of earlier reports would be to treat LD and UD as two actin dimers, one with tighter and other with open hydrodynamic shape which affects their migration patterns. While one of our models showed actins to be in face-to-face, other had actins in F-actin like edge-to-edge orientation. Former is compact globular, and the latter is relatively open, and would follow UD and LD like profile if cross-linked while bound to gelsolin. Furthermore, GA2 formed with face-to-face placed actins will lack nucleation ability as it will not allow binding of third or fourth actin, ternary complex having edge-to-edge orientation will be native-like seed for nucleation. Both arrangements are possible as per our SAXS data, and do not require large scale changes within complex. Additional arguments presented below may further present some possibilities to explain biochemical results reported on LD and UD bands.

Broader elution peak in SEC and Kratky plot of GA provided us first insight into the less globular nature of the binary complex of gelsolin and actin compared to GA2 complex. GA could also be formed via both depolymerization (even in F-buffer) and by binding to one unit of G-actin. Additionally, this GA complex failed to nucleate F-actin in G-buffer upon providing excess units of G-actin. Models of our binary complex suggested that the last actin unit or the first one is bound to C-terminal half of Ca^2+^-activated open gelsolin. Main difference between dummy residue model and residue details models were the possibility that G1 domain could be more open than that seen in the residue detailed models. Reason for this is because G1 domain is differentially positioned in unliganded vs. actin bound structures/shapes as it moves to latch on to actin bound to N-terminal half [2]. This observation provides an explanation for the much faster binding of the second actin molecule compared to the first one as it binds to the higher affinity site of G1. However, it is in disagreement with the partial proteolysis data which suggested that the first actin binds to the N-terminal half [14]. The latter conclusion was derived based on observed differences in the proteolytic profiles of the N-terminal half before and after the first actin binding. It is important to mention here that Ca^2+^-activated G1-G3 domains undergo more dramatic spatial repositioning than G4-G6 domains to bind respective actin units. G1 domains moves significantly via g1-g2 linker, while G2-G3 and G4-G6 remain more or less unchanged in 3D space from unliganded to actin bound shapes of Ca^2+^-gelsolin, except distal positioning by central g3-g4 linker [2]. This should lead to differences in partial proteolysis profiles of N-vs. C-terminal halves, and could provide skewed interpretation of whether actin is bound to N- or C-terminal half in GA. Our more probable model of GA generated by removing actin from N-terminal half suggests that to form GA2, second actin could come and bind N-terminal half. Closer of G1 domain on this second actin would position two actins in GA2 in face-to-face assembly seen in nucleation incompetent model. If GA must be generated from this GA2, detachment of G1 domain may dislodge actin bound to G2-G3 domains leading to C-terminal bound actin in GA. Considering the second model of GA, binding of second actin to N-terminal half would result in the two actins in anti-parallel orientation and would require remarkable “acrobatics” across g1-g2 and g3-g4 linker to position the two actins in parallel orientation. Though these possibilities provide easy explanations to LD and UD bands observed earlier, and reports on formation of either UD or LD in presence of ΔG1-gelsolin and G1-G3 half under nucleating conditions [17], they would suggest a simpler mechanism to form GA from GA2 as mentioned above. Formation of GA following a reversal across g3-g4 linker would require substantial disorder in the GA2 state which was clearly absent in the SAXS data. Few things are clear from our experiments and models, protein dynamics is not quenched post-formation of GA from GA2 or vice-versa. Even readjustments occur within GA and GA2 assemblies and they collectively regulate capping to seeding of nucleation of F-actin by Ca^2+^-gelsolin. While some understanding of the depolymerization mechanism could be made from earlier EM and SAXS work, significant improvements in our understanding of actin assembly by Ca^2+^-activated gelsolin from this work, like: 1) GA2 with edge-to-edge stacked actins like in F-actin could be the nucleation platform or seeding state, and 2) closely resembling GA2 with face-to-face stacked actins, and GA represent the capped or resting state of actin assembly.

## Acknowledgements

Funds from CSIR Network Projects UNSEEN and Bugs to Drugs were utilized for this study. AS and NP acknowledge research fellowships from CSIR and DBT INDIA. RG was a recipient of DST-WOSA Fellowship (SR/WOS-A/LS-220/2012). The authors are thankful to Dr. Cy Jeffries for his help in processing the SEC-SAXS data and the EMBL staff for help in SAXS data collection. Authors profoundly acknowledge support from Prof. JK Krueger and Dr. Matthew Paine for initial attempts to design some of these experiments when Ashish was in UNC Charlotte. Authors acknowledge consistent support of faculty and staff of IMTECH. This is IMTECH communication no. 143/2013.

## Additional Information

SASBDB entries of SAXS datasets available for open access are: <u>SASDUW3</u> [Calcium activated full-length gelsolin]; <u>SASDUX3</u> [F-actin in F-buffer at an actin concentration of 2 mg/mL]; <u>SASDUY3</u> [Calcium activated full-length gelsolin and the F-form of actin at a 1:32 molar ratio in F-actin buffer]; <u>SASDUZ3</u> [Calcium activated full-length gelsolin and the F-form of actin at a 1:16 molar ratio in F-actin buffer]; <u>SASDU24</u> [Calcium activated full-length gelsolin and the F-form of actin at a 1:8 molar ratio in F-actin buffer]; <u>SASDU34</u> [Calcium activated full-length gelsolin and the F-form of actin at a 1:4 molar ratio in F-actin buffer]; <u>SASDU44</u> [Calcium activated full-length gelsolin and the F-form of actin at a 1:2 molar ratio in F-actin buffer]; <u>SASDU54</u> [Calcium activated full-length gelsolin and the F-form of actin at a 1:1 molar ratio in F-actin buffer]; <u>SASDU64</u> [Calcium-gelsolin and the F-form of actin at a 1:2 molar ratio from high to low ionic strength]; <u>SASDU74</u> [Calcium-gelsolin and the F-form of actin at a 1:1 molar ratio from high to low ionic strength]; <u>SASDU84</u> [Calcium-gelsolin and the G-form of actin at a 1:2 molar ratio in low ionic strength (SEC-SAXS)]; <u>SASDU94</u> [Calcium-gelsolin and the G-form of actin at a 1:1 molar ratio in low ionic strength (SEC-SAXS)].

