## Supplementary Figures for "VISUALIZING THE NUCLEATING AND CAPPED STATES OF F-ACTIN BY Ca^2+^-GELSOLIN: SAXS DATA BASED STRUCTURES OF BINARY AND TERNARY COMPLEXES"

**Supplementary Figure S1:** SAXS intensity profiles of the basic reagents for experiments: G-actin in Buffer G, F-actin in F-buffer with fresh ATP and $Ca^{2+}$-gelsolin.

A

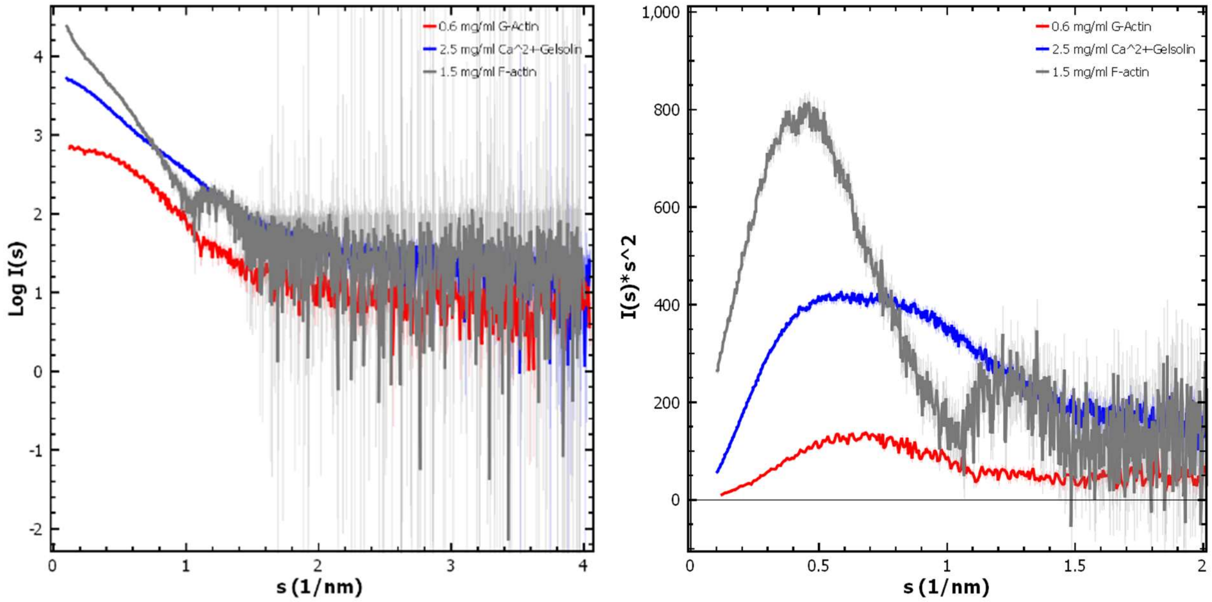


B

Guinier Approximation and Distance Distribution Profile for $Ca^{2+}$-gelsolin sample

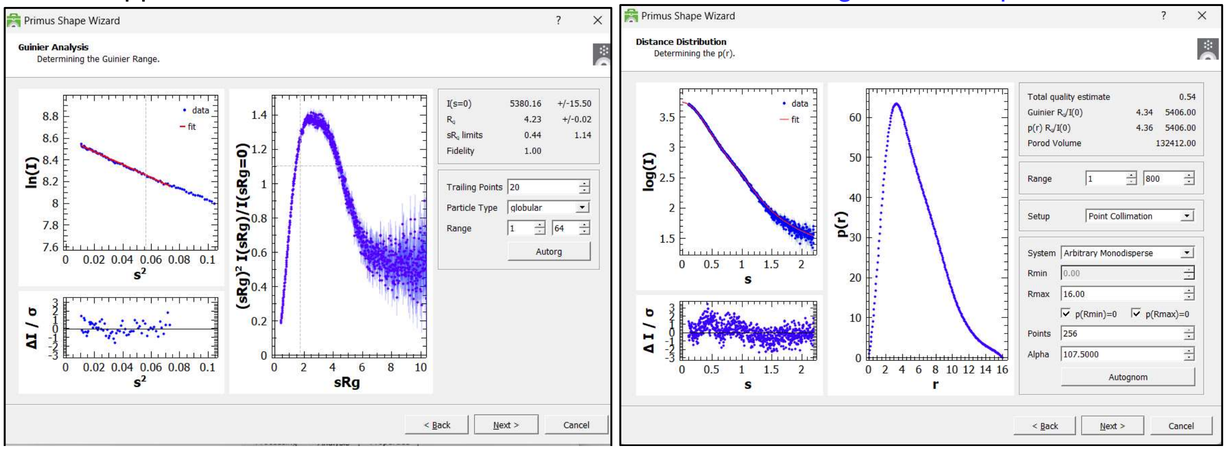

C

Guinier Approximation and Distance Distribution Profile for G-actin sample

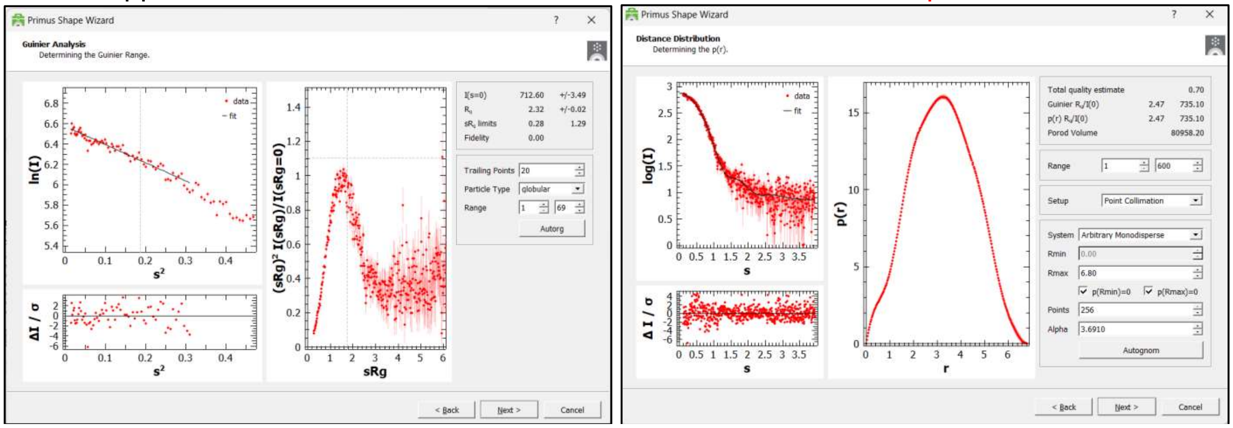


D

Guinier Approximation and Distance Distribution Profile for F-actin sample

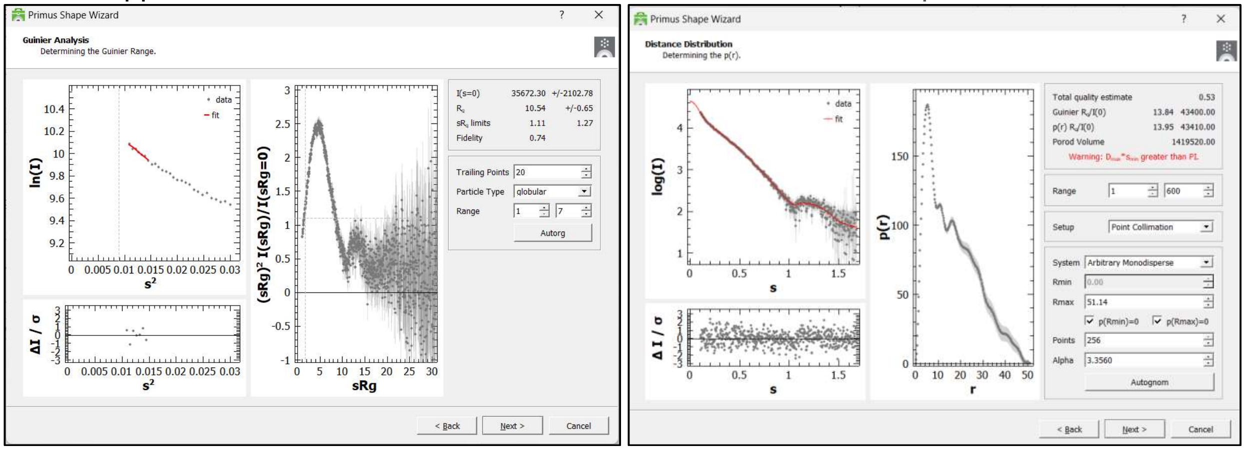


E

Comparison of Experimental SAXS profile with theoretical profiles of monomeric actin

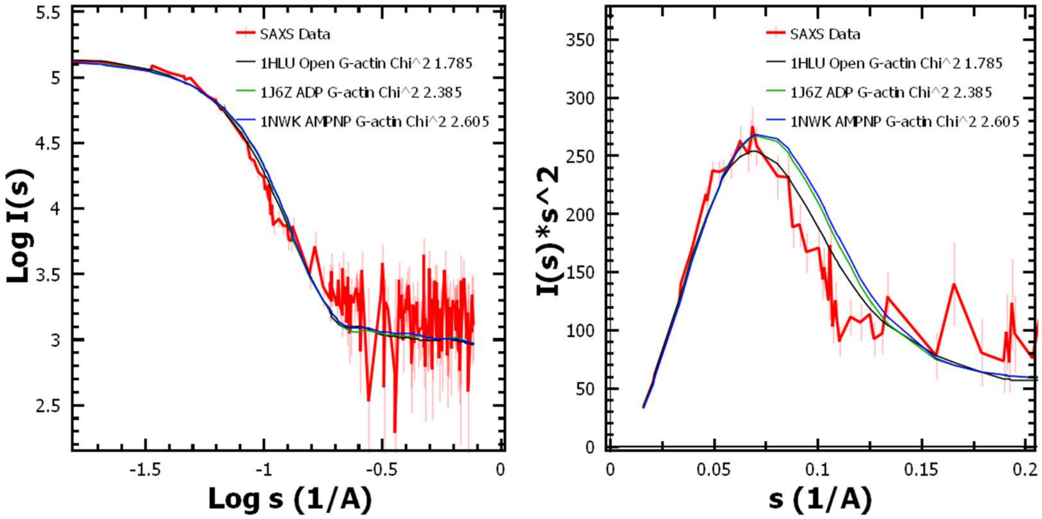

**Supplementary Figure S2:** SAXS intensity profiles of the F-actin samples at different concentrations in F-buffer are shown here in Double Log mode. Inset shows same datasets in Log-Linear mode devoid of the error bars to emphasize the depression in profiles around s~1 nm-1.

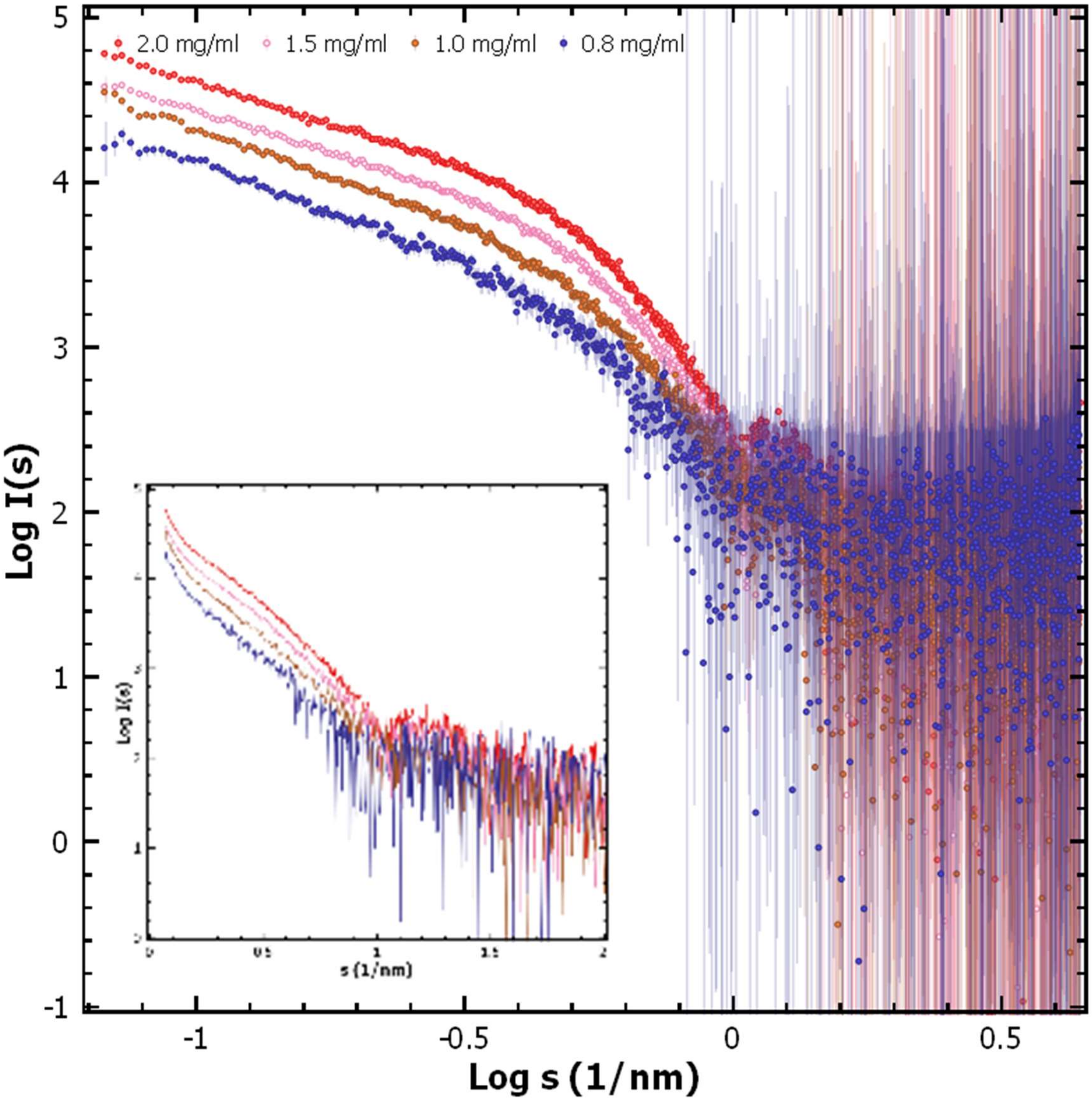

**Supplementary Figure S3:** (**A**) SAXS intensity profile of $Ca^{2+}$-gelsolin at 2 mg/ml is shown in Double Log mode. Inset shows the same profile in Log-Linear mode. (**B**) Computed SAXS profiles from the structure of inactive gelsolin (PDB ID 1D0N Chain A) and constructed model of $Ca^{2+}$-activated open structure are compared with the experimental data. Legends show the color of the plotted curves and computed $\chi^2$ values. (**C**) Two rotated views of the model of fully active or open structure of gelsolin after stapling the G1-G3 and G4-G6 structures to the model of g3-g4 linker. (**D**) SAXS profiles computed for new models computed for the open structure(s) of open gelsolin. The initial profile is shown by black lines. $\chi^2$ values of the comparison are mentioned in the legend of **Figure 1B**.

A

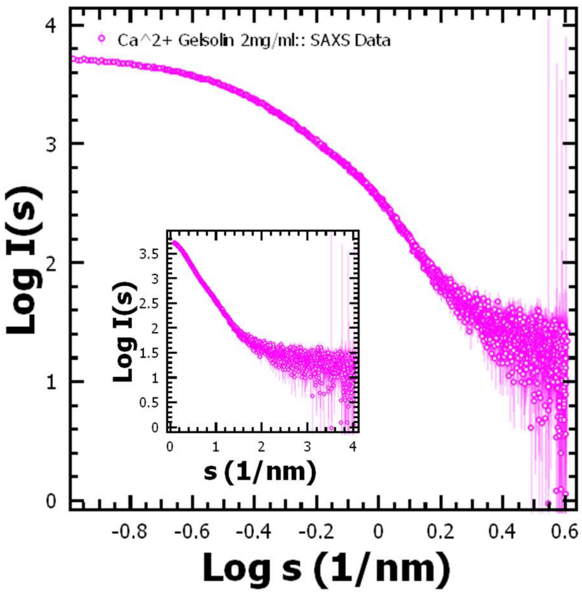


B

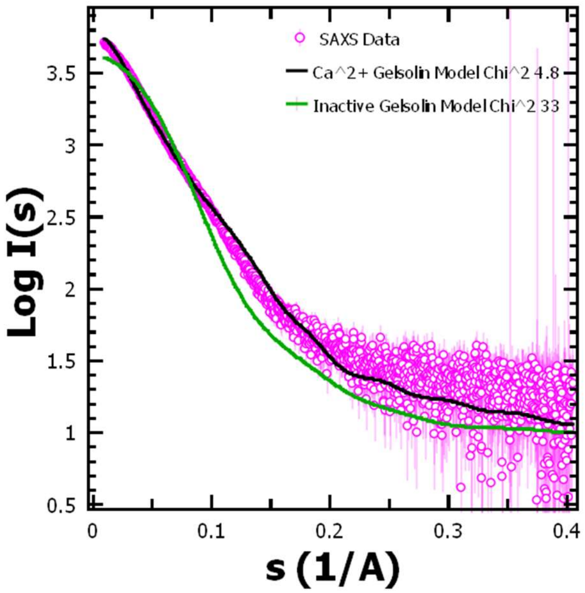


C

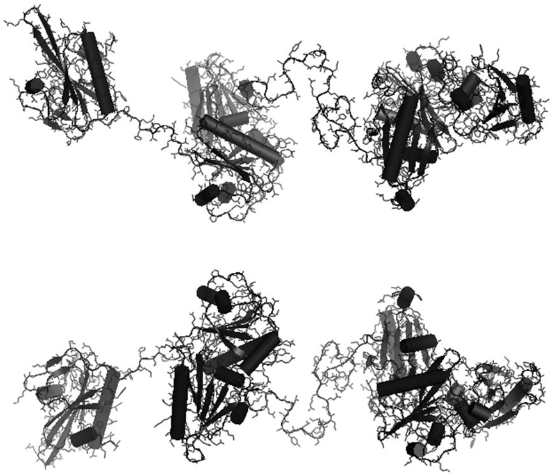

D

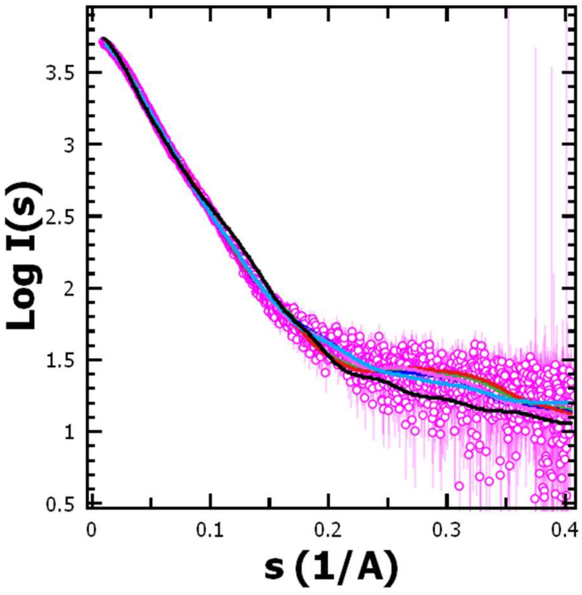

**Supplementary Figure S4:** (**A**) SAXS profiles of different molar mixtures of actin in F-form and $Ca^{2+}$-gelsolin are shown in Double Log mode. (*Repeated Figure 1D here for direct comparison*). (**B**) Same ratios as in panel A, but the gelsolin sample lacked free calcium. (**C**) Guinier plots for rod shape i.e. ln I(s)*s vs. $s^2$ have been shown for four F-actin: $Ca^{2+}$-gelsolin mixtures have been shown here, The black lines are the linear fit to negative slope data points. The ratios are mentioned in the right corner of the plots. Lower plots are respective fits to the linear region.

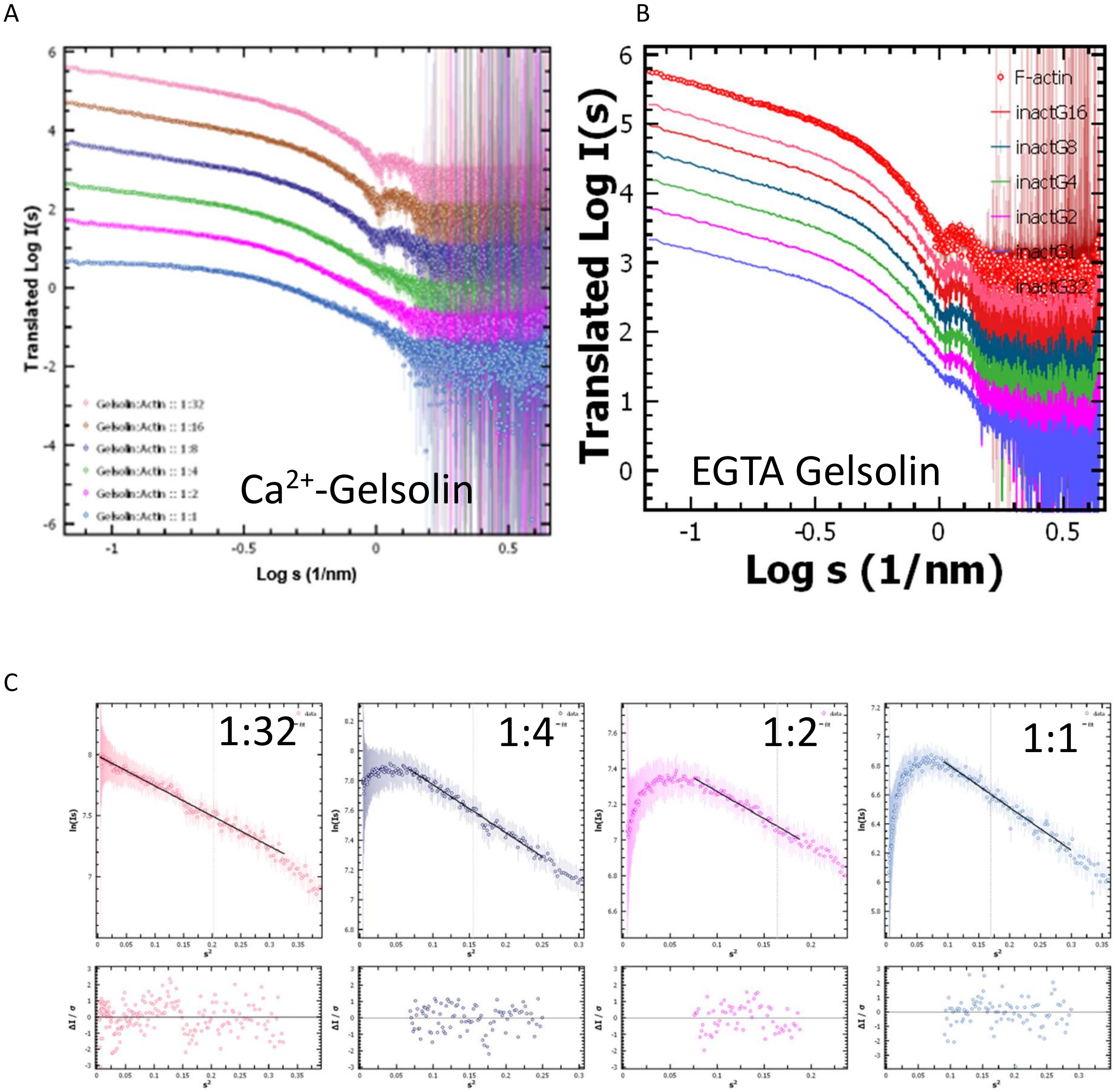

**Supplementary Figure S5: (A)** SAXS profiles of mixtures at different time points of incubation. The mixtures were GA2 (left) and GA (right) entity prepared in G-buffer and then mixed with G-actin and then dialyzed against G- or F-buffer. The profiles are of mixtures dialyzed against G-buffer (**upper panels**) and against F-buffer (**lower panels**). (**B**) $R_g$ values estimated by AutoRg program for different datasets shown in panel A are plotted here as a function of incubation time.

A

GA2 + G-actin in G-buffer
Then, dialyzed vs. G-buffer

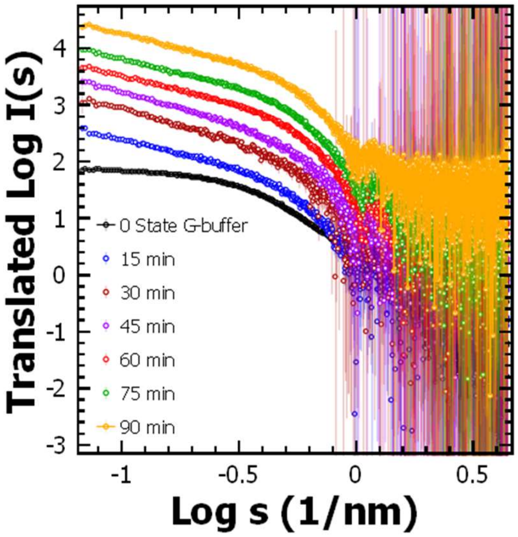


GA + G-actin in G-buffer
Then, dialyzed vs. G-buffer

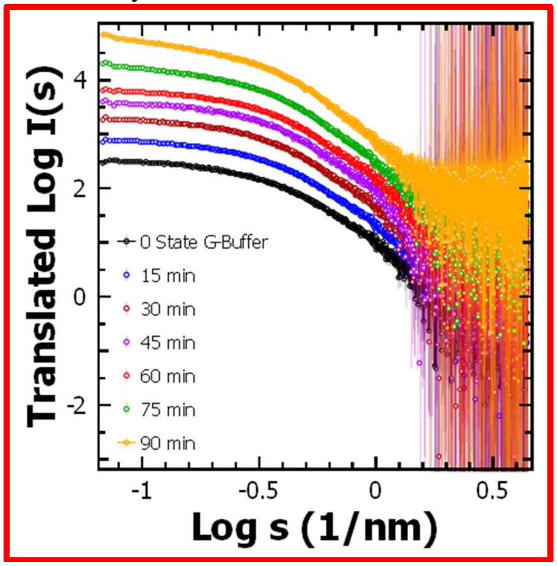


GA2 + G-actin in G-buffer
Then, dialyzed vs. F-buffer

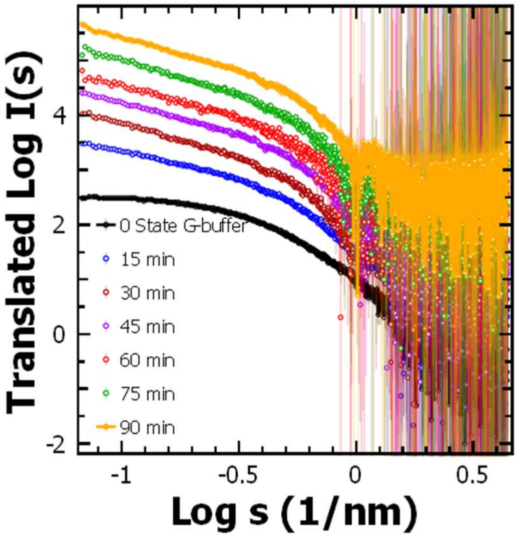


GA + G-actin in G-buffer
Then, dialyzed vs. F-buffer

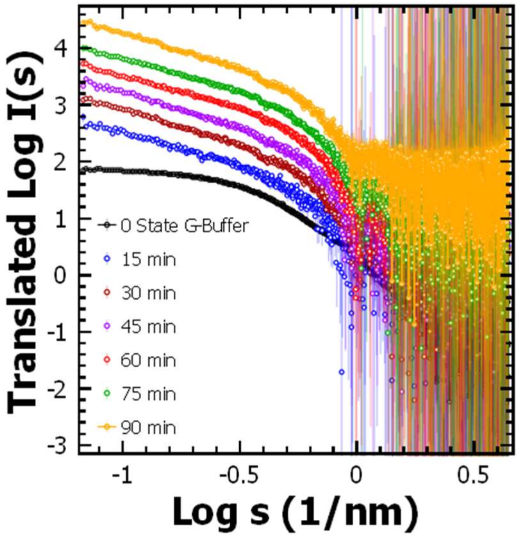

**B**

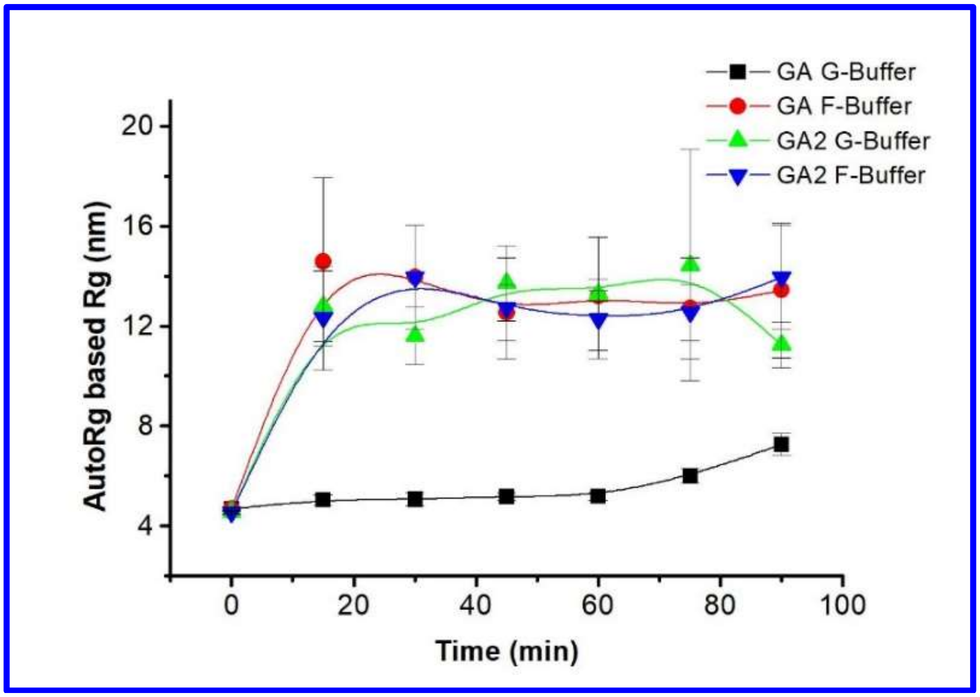

GA G-Buffer
GA F-Buffer
GA2 G-Buffer
GA2 F-Buffer
AutoRg based Rg (nm)
20
16
12
8
4
0
20
40
60
80
100
Time (min)

**Supplementary Figure S6:** SAXS data analysis of the two SAXS datasets generated by averaging data points eluting from SEC-SAXS of 1:2 (A) and 1:1 (B) molar mixtures of $Ca^{2+}$-gelsolin and G-actin in G-buffer. The left panels in upper row shows the Guinier approximation to the data using globular shape profile. The black line shows the linear fit to the points in the plot. Other plots are the fit and normalized Kratky plots. The right panel is a zoomed in image of the peak height of the normalized Kratky plot. Lower panels are the distance distribution profiles and parameters deduced by this approximation. (C) Snapshots of the molecular masses estimated for the predominant species in the two datasets.

A

SAXS analysis of 1:2 mixture from mixing $Ca^{2+}$-gelsolin and G-actin followed by SEC-SAXS

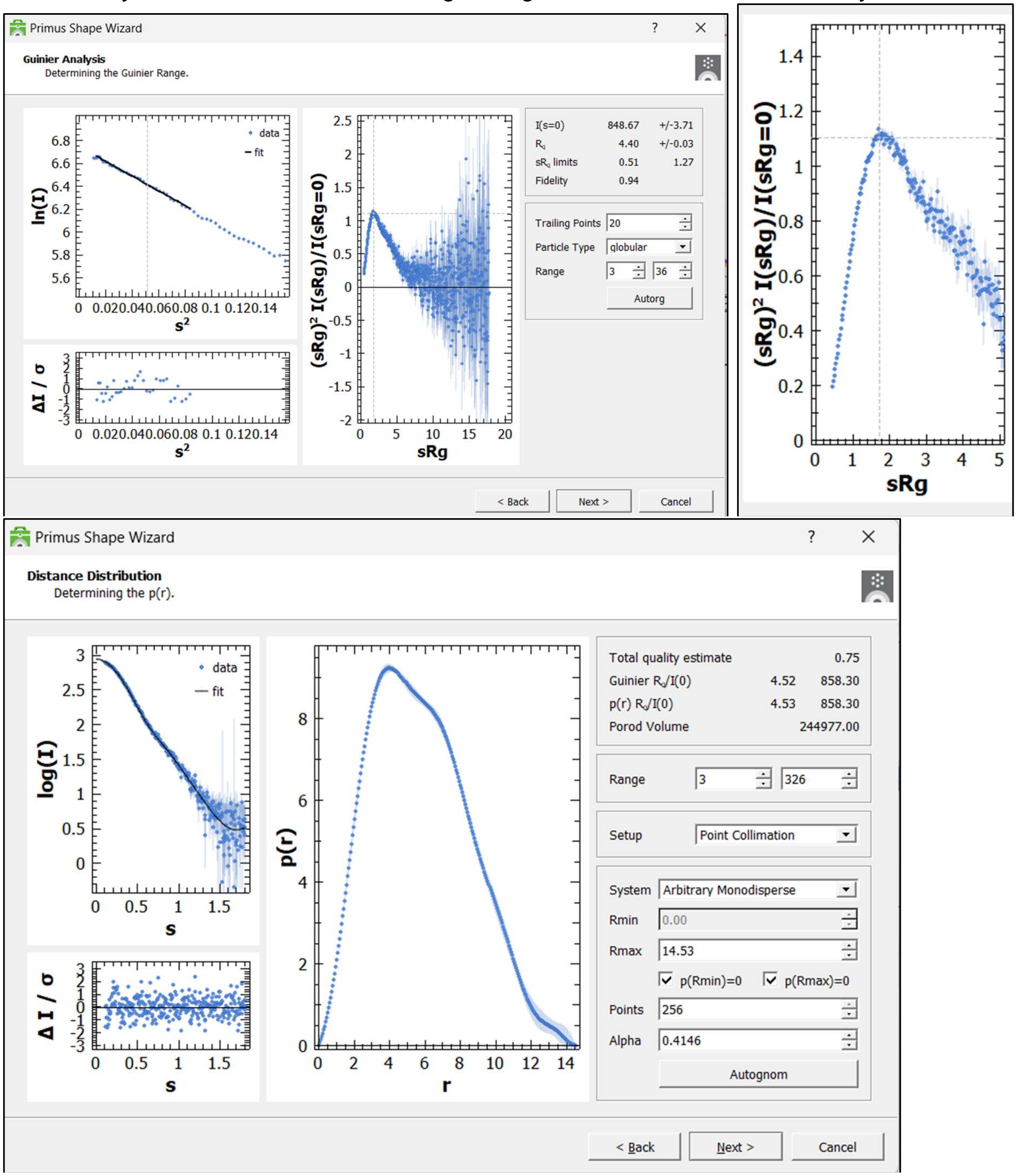

**B**

SAXS analysis of 1:1 mixture from mixing $Ca^{2+}$-gelsolin and G-actin followed by SEC-SAXS

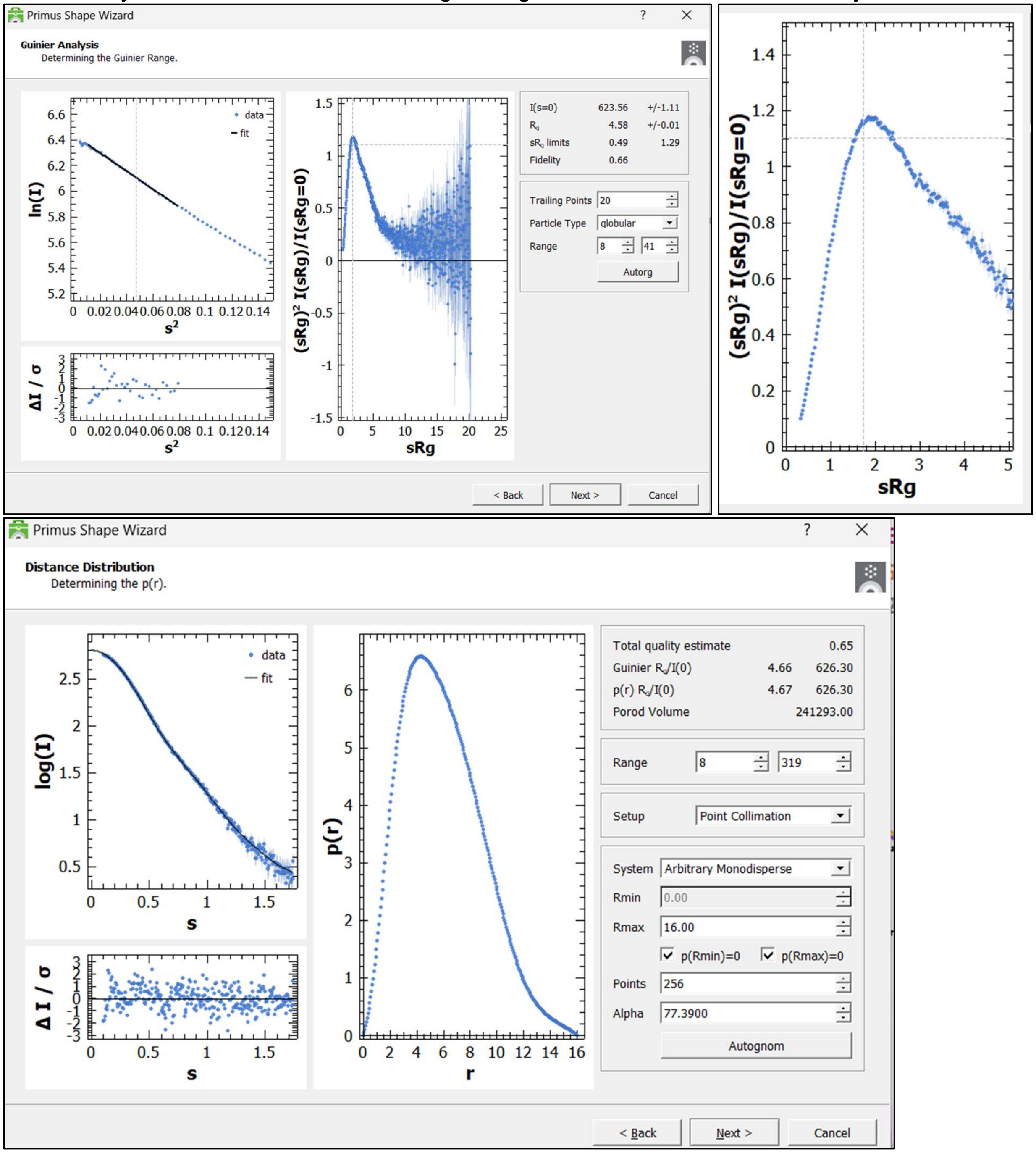

C

Molecular weight estimation of scattering species in 1:2 (left) and 1:1 (right) mixtures of $Ca^{2+}$-gelsolin and G-actin followed by SEC-SAXS

1:2 Mixture

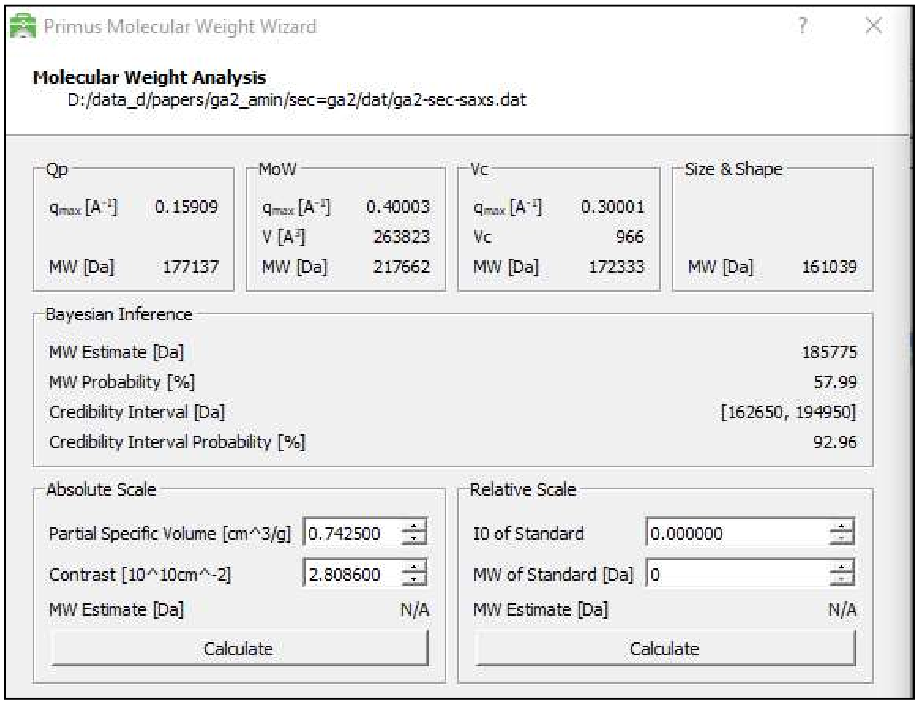


1:1 Mixture

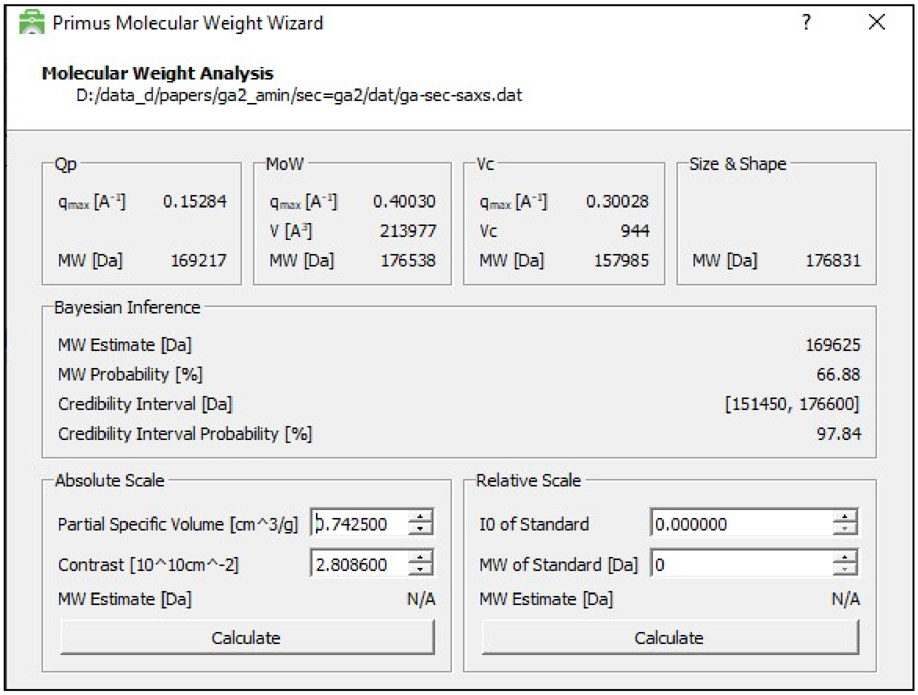

**Figure S7:** Images **and details** of the chain-ensemble model solved for GA2 complex from its SAXS dataset and its superimpositions with the residue models generated as described in the manuscript. For each set, three rotated views have been shown.

A GASBOR Model with $\chi^2$ of 0.76 to SEC SAXS Data of GA2 mixture (dummy Asp and water molecules are shown as violet and gray cpk, respectively)

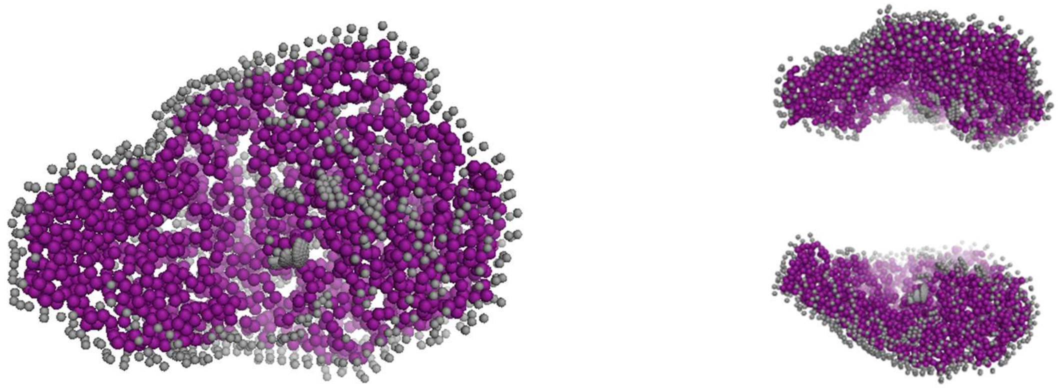

B Model 8 of ternary model (Blue lines) with $\chi^2$ of 0.70 to SEC SAXS Data of GA2 mixture

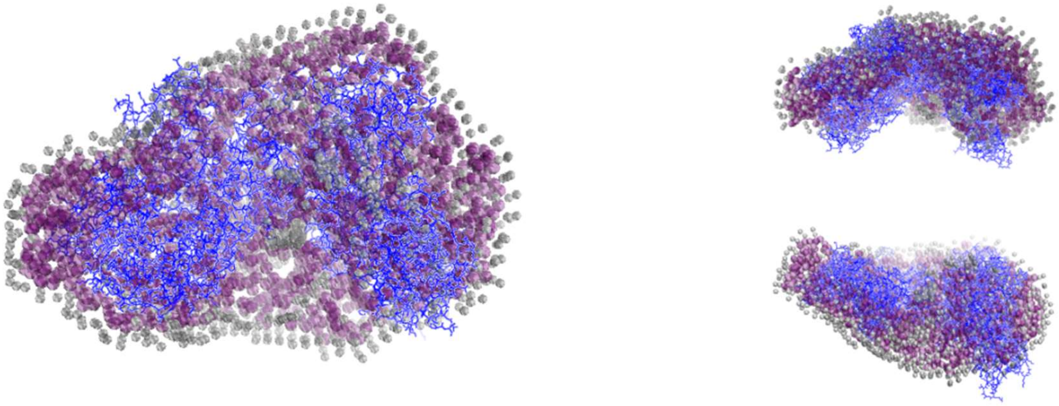

C Model 2 or 6 of ternary model (Red lines) with $\chi^2$ of 1.15 to SEC SAXS Data of GA2 mixture

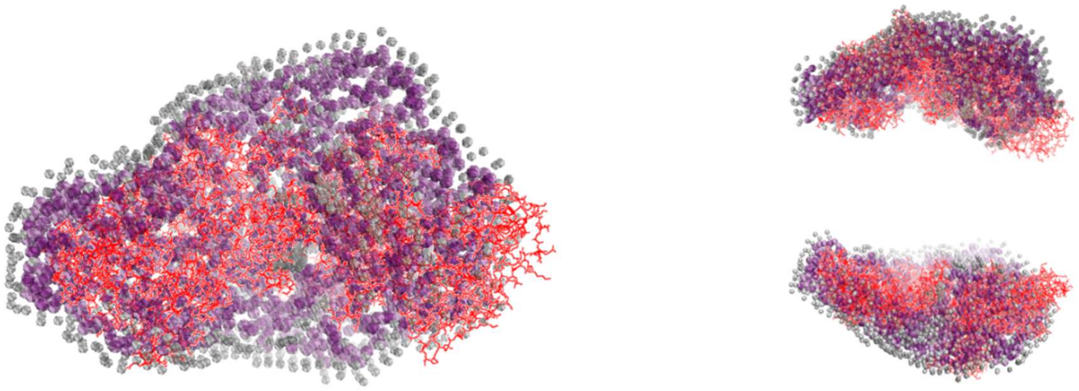

**Figure S8:** Images **and details** of the chain-ensemble model solved for GA complex from its SAXS dataset and its superimpositions with the residue models generated as described in the manuscript. For each set, three rotated views have been shown.

A GASBOR Model with $\chi^2$ of 0.82 to SEC SAXS Data of GA mixture (dummy Asp and water molecules are shown as orange and gray cpk, respectively)

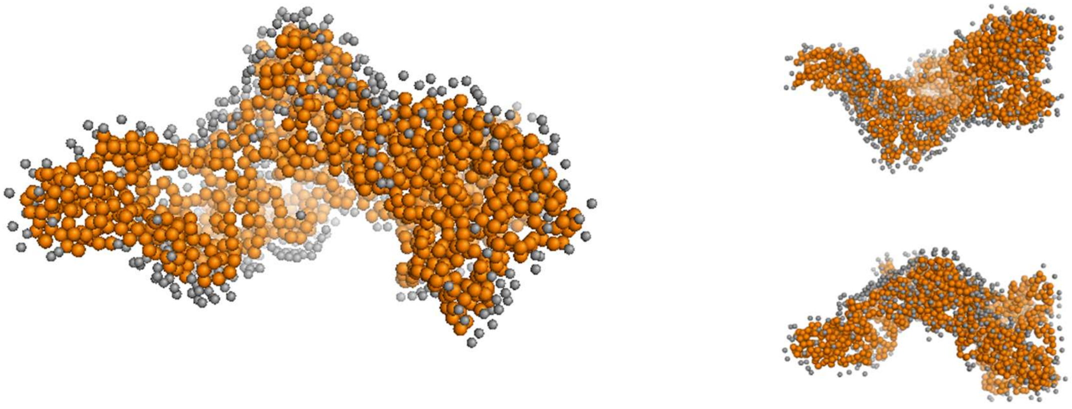

B Model 3 of binary model (Magenta lines) with $\chi^2$ of 6.68 to SEC SAXS Data of GA mixture

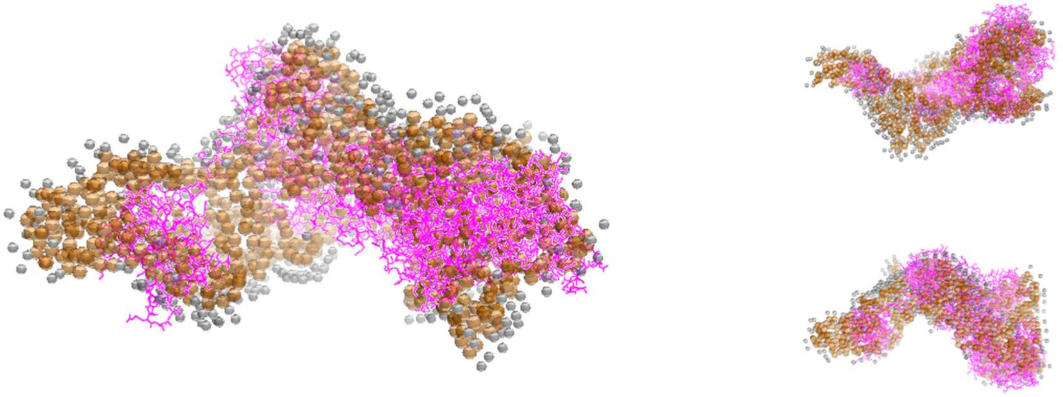

C Model 1 of binary model (Forrest green lines) with $\chi^2$ of 10.31 to SEC SAXS Data of GA mixture

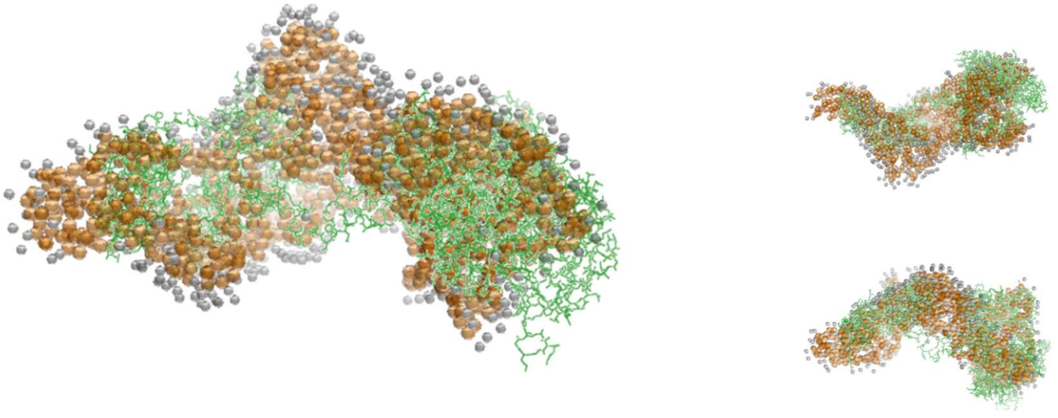
